# Extracting informative glycan-specific ions from glycopeptide MS/MS spectra with GlyCounter

**DOI:** 10.1101/2025.03.24.645139

**Authors:** Kathryn Kothlow, Haley M. Schramm, Kayla A. Markuson, Jacob H. Russell, Emmajay Sutherland, Tim S. Veth, Ruby Zhang, Anna G. Duboff, Vishnu R. Tejus, Leah E. McDermott, Laura S. Dräger, Nicholas M. Riley

**Affiliations:** Department of Chemistry, University of Washington, Seattle, WA, 98195

**Keywords:** Glycoproteomics, informatics, glycopeptides, tandem mass spectrometry, data evaluation

## Abstract

Glycopeptide tandem mass spectra typically contain numerous glycan-specific fragments that can inform several features of glycan modifications, including glycan class, composition, and structure. While these fragment ions are often straightforward to observe by eye, few tools exist to systemically explore these common glycopeptide spectral features or explore their relationships to each other. Instead, most studies rely on manual inspection to understand glycan-informative ion content in their data, or they are restricted to evaluating the presence of these ions only in the small fraction of spectra that are identified by glycopeptide search algorithms. Here we introduce GlyCounter as a freely available, open-source tool to rapidly extract oxonium, Y-type, and custom ion information from raw data files. We highlight GlyCounter’s utility by evaluating glycan-specific fragments in a diverse selection of publicly available datasets to demonstrate how others in the field can make immediate use of this software. In several cases, we show how conclusions drawn in these publications are evident simply through GlyCounter’s extracted ion information without requiring database searches or experiment-specific programs. Although one of our goals is to decouple spectral evaluation from glycopeptide identification, we also show that evaluating oxonium ion content with GlyCounter can supplement a database search as valuable spectral evidence to validate conclusions. In all, we present GlyCounter as a user-friendly platform that can be easily incorporated into most glycoproteomic workflows to refine sample preparation, data acquisition, and post-acquisition identification methods through straightforward evaluation of the glycan content of glycoproteomic data. Software and instructions are available at https://github.com/riley-research/GlyCounter.

## INTRODUCTION

Glycosylation is an important and prevalent protein modification implicated in numerous biological processes that govern health and disease. Indeed, the better our tools to study glycosylation have become, the more diseases we find with altered glycosylation as critical molecular features.^1–4^ Mass spectrometry (MS) has advanced to play a central role in the study of glycoproteins, and glycopeptide sequencing by tandem MS (MS/MS) is the gold standard for site-specific glycoprotein analysis. Collisions, electrons, and photons have all been successfully employed for glycopeptide MS/MS.^5^ Given the labile nature of glycosidic bonds, many MS/MS methods fragment the glycan itself, but sequence-informative fragments derived from the peptide backbone can also be ubiquitous.^6^ The multiple glycan- and peptide-specific fragmentation modalities possible in a given glycopeptide precursor ion typically lead to complex MS/MS spectra that remain an informatic challenge in glycoproteomics.^7–9^ Despite this complexity, glycan-specific fragmentation commonly produces two ion types that are hallmarks of glycopeptide spectra and are crucial for accurate glycan characterization. B-ions (often referred to as oxonium ions, albeit in the strictest sense not all oxonium ions are B-ions) consist of “glycan-only” fragments from the terminal (i.e., non-reducing) end of the glycan that do not contain the peptide backbone. Conversely, Y-ions comprise the peptide backbone with a truncated form of the still-attached glycan that is missing a piece(s) of the terminal end. Critically, glycan-specific fragments serve as diagnostic ions that 1) indicate whether a spectrum represents a glycan-containing molecule, 2) provide information on glycan composition, and 3) reveal structural details such as monosaccharide linkage information.^10–33^

Software tools that interpret glycopeptide MS/MS spectra have advanced dramatically, especially in recent years^34^, and most modern glycoproteomics search tools incorporate glycan-specific ions, especially oxonium ions, into scoring algorithms when identifying spectra (e.g., MSFraggerGlyco^35^, Byonic^36^, pGlyco3^37^, O-Pair Search^38^, GlycoDecipher^39^, StrucGP^40^). Access to the oxonium ion information, however, is limited to identified spectra, leaving any glycan-specific fragments in unassigned spectra unannotated and often uninvestigated further in data processing. Because overlap between search algorithms is poor, this exclusion of data means that potentially useful spectra are unwarrantedly discarded. Analysts may also remain blind to useful data hiding in plain sight in these situations, too, because they unknowingly used suboptimal or incorrect search parameters. Omitting these spectra becomes an unforced error in data processing simply because tools to make informed decisions are not available. This issue becomes even more glaring when considering that glycopeptide identifications are often a metric used for method development. If informative spectra remain unused in method assessment, proper evaluation is difficult to achieve. Instead, we argue that many of these situations can be resolved by simply decoupling glycan-specific ion evaluation from glycopeptide identification and using glycan-specific ions as metrics to rapidly interrogate data and inform downstream analysis decisions.

Inspecting raw data for glycan-specific ions differs by MS vendor, but filtering for user-defined mass-to-charge (m/z) values can commonly be done with tools like Freestyle and MSConvert^41^. While useful for observing glycan-specific ions, these approaches do not directly report the number, co-occurrence, intensity, or other metrics for user-defined ions, requiring additional time-consuming steps to obtain these data or investigate shared patterns between multiple ions. Simply put, these tools make glycan-specific ions easy to observe, but difficult to extract. ProteinProspector has a built-in feature called MS-Filter that can filter MS/MS scans for the presence of user-defined glycan-specific fragment ions.^42,43^ However, ProteinProspector returns a filtered raw file for subsequent use in a glycopeptide search engine, meaning glycan-specific fragment ion data must still be extracted. pGlyco3 provides a Python script for extracting B- and Y-ions (https://github.com/pFindStudio/pGlyco3/tree/main/scripts/BY_ion_extractor.py) which does not contain a user interface, limiting accessibility. It also requires the user to enter lists of B-ions and Y-ions to search for by hand. Given this, we found ourselves wanting a more flexible and straightforward tool to quantitatively extract glycan-specific ion intensity from raw data to expand access to the information they encode.

Here we introduce GlyCounter, a simple, freely available, open-access tool written in C# that extracts glycan-specific fragment ion information from raw data files. GlyCounter allows for the upload of Thermo .raw and .mzML files, comes with over 50 predefined common glycan-specific fragment ions, and has the option to upload user-defined custom ions. GlyCounter has been released as an open-source Windows Form .NET application available at https://github.com/riley-research/GlyCounter. To demonstrate its functionality and utility in glycopeptide workflows, we present multiple analyses of published, publicly available data using GlyCounter and highlight its versatility for many applications across glycoproteomics.

## EXPERIMENTAL PROCEDURES

### Development and processing approach

GlyCounter was developed as a C# Windows Form application in .NET 8.0. The application has been released as an open-source repository available at https://github.com/riley-research/GlyCounter. A standalone executable (GlyCounter.exe) is available in the Releases section of the GitHub repository. GlyCounter runs through a straightforward graphical user interface (GUI) and first creates a hash set of oxonium ions, including any custom ions input by the user. Then for each scan at the user-defined MS levels, GlyCounter reads the dissociation type, total ion current (TIC), and peak information before checking the experimental spectrum for each m/z value from the hash set. Peaks are filtered by either signal-to-noise or an intensity threshold depending on the file type, with signal-to-noise being the preferred choice if provided by the raw data. If an oxonium ion is found within a user-defined tolerance (a default of 15 ppm was used in all analyses here unless otherwise stated), its peak depth (i.e., its relative rank based on intensity) and ion intensity are recorded. Fragment ppm tolerance is instrument dependent, as both resolution and calibration metrics can have effects on how close an ion’s experimental mass-to-charge value is to its theoretical counterpart. 15 ppm was chosen as the default as this will enable straightforward use with many of the settings used in typical glycoproteomics experiments (e.g., 30k resolution at m/z 200 with Orbitrap mass analysis), but this can be easily tailored by the user to the mass accuracy available for a given experiment. The settings also allow for a Dalton fragment tolerance if the user is working with low resolution data, e.g., when an ion trap is used for mass analysis. GlyCounter outputs three tab-delimited text files to display oxonium ion peak depth, signal, and a summary of scan types and oxonium ion content (**Supplementary File 1**). For each spectrum, GlyCounter assigns a Boolean “LikelyGlycoSpectrum” classifier based on the number of oxonium ions found, their percentage contribution to the TIC, and their peak depth. For number of oxonium ions found in HCD and UVPD spectra, the default requirement to be considered LikelyGlyco scales with number of oxonium ions selected by the user: 4 oxonium ions are required for fewer than 6 oxonium ions checked, half the number of selected ions are required for 6 through 15 oxonium ions selected, and 8 oxonium ions are required if more than 15 ions are selected. To be considered LikelyGlyco, at least 4 oxonium ions need to be matched. If a user selects fewer than 4 ions, GlyCounter will extract information about them, but no spectrum will be marked LikelyGlyco. These values are halved for ETD spectra, where ETD here and throughout refers to any ETD-centric dissociation type including AI-ETD, EThcD, and ETcaD. For peak depth settings, oxonium ions that count toward the thresholds above must be found within the 25, 50, and 25 most abundant peaks for HCD, ETD, and UVPD spectra, respectively. Default contribution of oxonium ions to TIC is set to fractions of 0.20, 0.05, and 0.20 (i.e., 20%, 5%, and 20%) of TIC for HCD, ETD, and UVPD, respectively. Default settings were assigned through empirical evaluation provided in **Supplementary Figures 1-6**, which appears robust when comparing with a true null *E. coli* sample that does not contain glycopeptides (**Supplementary Figure 7**). Note, extensive UVPD data was not available, but testing showed that HCD and UVPD spectra had similar qualities for oxonium ion abundances. With the common ions box selected, the default settings were as follows: For higher-energy collisional dissociation (HCD) or ultraviolet photodissociation (UVPD) scans, 8 of the 25 most abundant peaks must be oxonium ions, and the sum of all oxonium ions must be at least 20% of the TIC. For electron transfer dissociation (ETD), 4 of the 50 most intense peaks must be oxonium ions, and all oxonium ion signal must sum to at least 5% of the TIC. These default settings were used for all GlyCounter searches unless otherwise stated, and all data here was taken directly from GlyCounter outputs and plotted in R. Even though these are default settings, users have access to change all of them through the GUI. The search speed of GlyCounter was compared directly to an MSFragger-Glyco search of the same raw file and found to be approximately three times faster (**Supplementary Figure 8**). **Figure 1** displays the GlyCounter user interface, and a tutorial that describes step-by-step instructions for using the program is available in **Supplementary File 2**. With many options for customizability, GlyCounter provides a user-friendly platform for extracting ions from mass spectrometry data.

**Figure 1.**
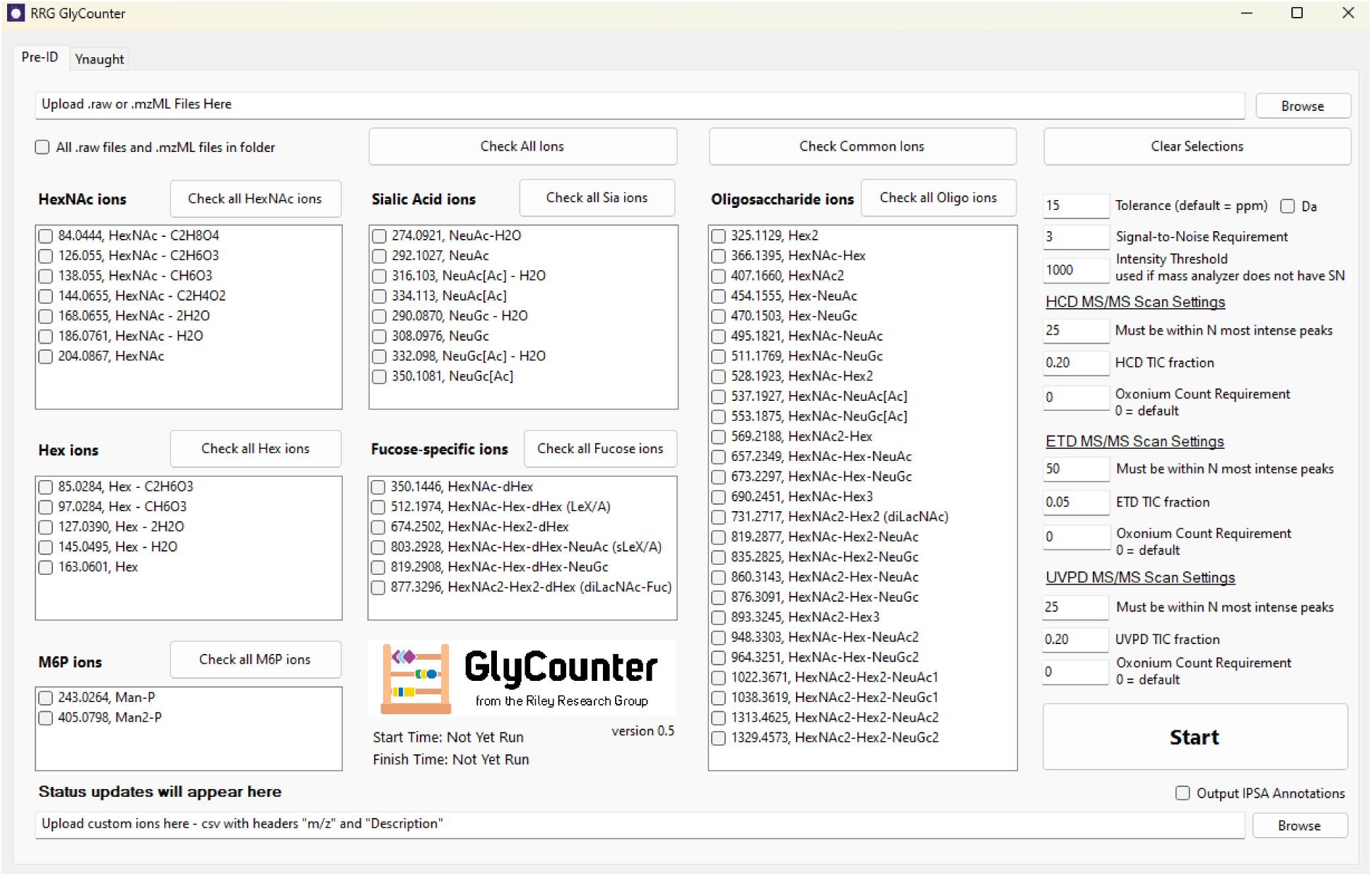
The GlyCounter interface. The user interface of GlyCounter allows the user to input an MS .raw or .mzML file and any custom ions to consider. The 50+ default ions include oxonium ions from several monosaccharides and oligosaccharides that can be selected individually or in groups. The “Check Common Ions” feature includes oxonium ions that are typically considered in our group and in studies across the field. Several user-defined parameters enable selection of mass tolerance for ion detection in ppm or Da, signal-to-noise threshold for considering ions as detected and settings for multiple dissociation types that define the “Likely Glycopeptide” classification assigned by GlyCounter for spectra with specified features. Raw data files can be run individually or batched for all raw data in a given folder location. Additionally, the Ynaught module, shown in Supplementary Figure 9, allows users to extract Y-type ions from spectra assigned glycopeptide identifications.

### Input requirements

GlyCounter is compatible with both Thermo .raw and .mzML spectral file formats. MSConvert can be used to convert many file formats to .mzML, and GlyCounter can read any MS/MS scan fragmented with HCD, ETD, EThcD, or UVPD. GlyCounter requires that .mzML files not be zlib compressed or packaged in gzip. MS/MS scans must contain only centroided peaks, and the conversion from profile to centroid can also be done in MSConvert by selecting the Peak Picking filter and including MS level 2. Files can be processed either one at a time or in batches. Input spectral files are read using the Nova package^44^ (https://schweppelab.github.io/Nova/), and spectra are processed with a combination of custom code and the C# Mass Spectrometry Library (https://github.com/dbrademan/CSMSL).

### The Ynaught Module

With the release of GlyCounter, we are also introducing a supplementary module built into GlyCounter to easily identify Y-type ions called Ynaught. The Ynaught interface is shown in **Supplementary Figure 9**. Ynaught supports both Thermo .raw file and .mzML file types as an input. Instead of being independent of the search results, Ynaught uses an MSFragger-Glyco search output from FragPipe to find identified glycopeptides in the spectra. Search files from other search engines need to be formatted to match the output from FragPipe using the example file in the repository as a guide. In addition to uploading the search results and a raw file, Ynaught requires a glycan database that has the glycan names/compositions and masses used in the database search. The user can pick which Y-ions to search for, as well as assign a ppm tolerance value and signal-to-noise threshold. Ynaught has two different ways to define Y-ions. The first is the monosaccharide composition remaining on the peptide after fragmentation. The second is the ion resulting from a neutral loss of monosaccharides from the precursor glycopeptide. To calculate the m/z values of these ions, Ynaught uses the mass of the identified peptide backbone with no glycan modification and the precursor glycopeptide, respectively. Ynaught also can look for isotope peaks when searching for Y-ions, including M+1 and M+2, which are selected by user choice. Larger glycopeptide fragments have the potential to be at any charge state between +1 and the precursor charge state. The charge state limits in Ynaught are set based on the precursor charge “P” and can be integer values or references to P (e.g., from charge = 1 to charge = P-1). Ynaught offers two optional uploads for custom masses to add to the unmodified peptide mass and subtract from the intact precursor mass. Ynaught outputs files with the Y-ion peak depth, Y-ion signal, and a summary of the results in similar formats to oxonium ion outputs from GlyCounter.

### Analyzing Publicly Available Data

We searched several publicly available MS datasets with GlyCounter to demonstrate the use cases and highlight the value GlyCounter adds to 1) view oxonium ions outside the context of a database search and 2) investigate patterns of co-occurring fragment ions, all file type conversion and peak picking was done using MSConvert. The data used are accessible at ProteomeXchange Consortium dataset identifiers PXD023448, PXD011533, PXD001571, PXD035775, PXD001404, PXD005655, PXD010333, PXD022988, PXD005411, PXD005413, PXD005412, PXD005553, PXD005555, PXD041217, PXD023448, PXD004559 and MassIVE repositories MSV000091172, MSV000094544, MSV000083070.

## RESULTS

### Examining glycopeptide fragmentation

Different dissociation methods used in glycoproteomics experiments reveal unique information about glycopeptides. Due to glycan heterogeneity, these informative spectral features are integral to identifying and localizing glycopeptides. B- and Y-type ions are examples of such spectral features, in many cases providing sufficient evidence to assign glycan compositions. To better understand how MS/MS parameters contribute to glycan-specific ion variability, we used GlyCounter to compare several common and more bespoke dissociation types used for glycoproteomics. First, we re-evaluated a dataset that compared higher energy collisional dissociation (HCD), stepped collision energy HCD (sceHCD), electron transfer dissociation (ETD), and ETD with supplemental HCD activation (EThcD) for N-glycopeptide fragmentation.^45^ Even in this large dataset, GlyCounter makes it easy to extract oxonium ion signals for each method (**Figure 2A**). These data show that even though total oxonium ion signal may peak at higher HCD energies, specific oxonium ions that may be helpful for compositional or structural assignment have their highest signal with EThcD, low HCD energies, and sceHCD. Considering this, we also wanted to show how GlyCounter can help the recent movement toward structural glycoproteomics.^46^ Maliepaard et al. recently used HCD collision energy ramps to generate “breakdown curves” that can reveal structural information between glycopeptides harboring different glycan isomers.^47^ Their data were made by custom scripts tailored specifically to their study, so we wanted to highlight the flexibility of GlyCounter as a tool to rapidly recreate their charts (**Figure 2B**). We were able to generate effectively identical figures in just minutes. Our recreations display GlyCounter’s ability to eliminate the need for experiment-specific programs for extracting these ion types, making structural glycoproteomics more accessible for future studies.

**Figure 2.**
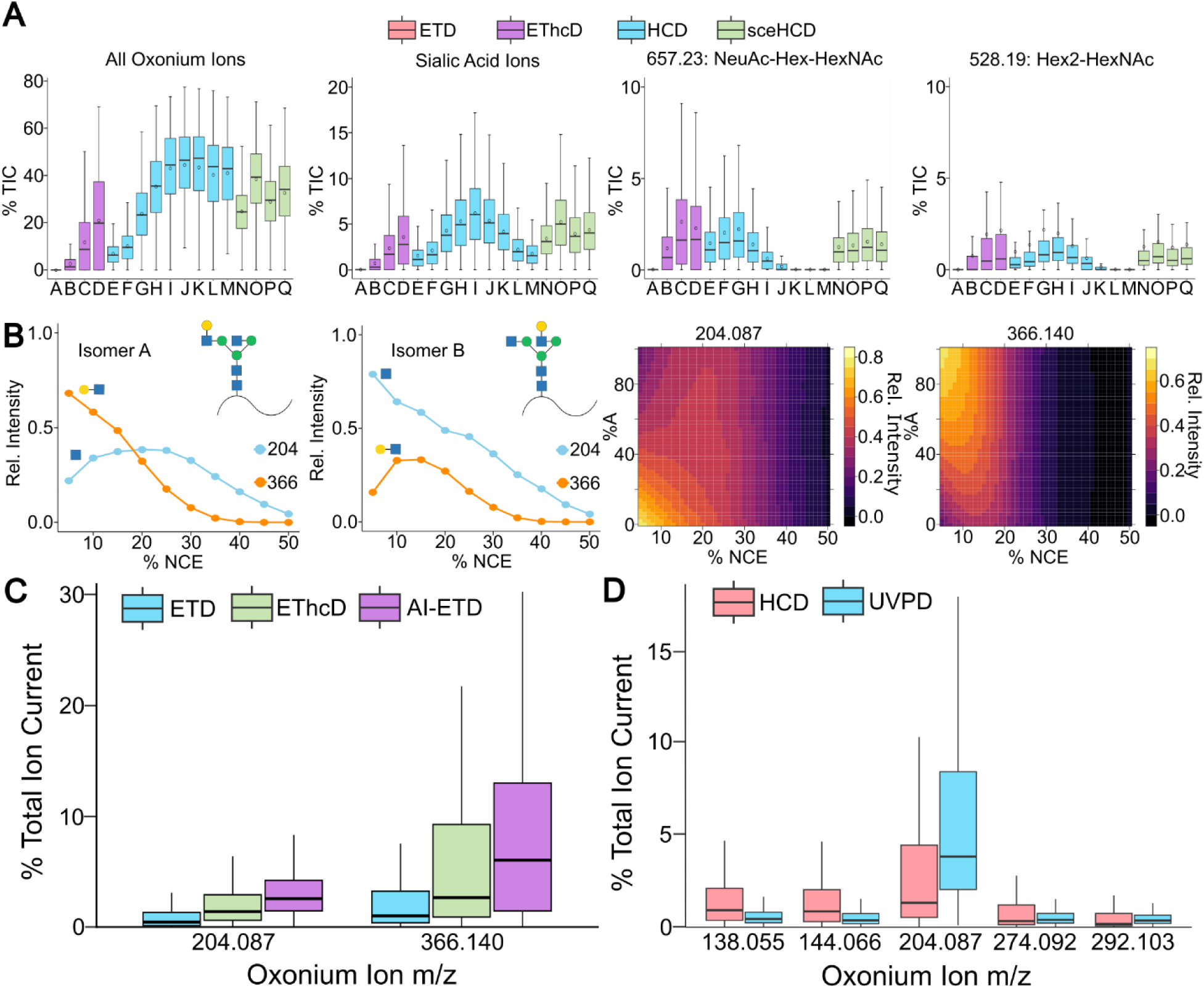
Analyzing glycopeptide dissociation types with GlyCounter. **A)** Boxplots show the percent total ion current (TIC) for all oxonium ions (far left), sialic acid ions m/z 274 and 292 (middle left), NeuAc-Hex-HexNAc ion (middle right), and Hex2-HexNAc ion (far right). The dissociation methods used have been represented by letters. A: ETD, B: EThcD with HCD set to 15 normalized collision energy (NCE), C: EThcD with 25 NCE, D: EThcD with 35 NCE, E: HCD at 10 NCE, F: HCD at 12 NCE, G: HCD at 20 NCE, H: HCD at 25 NCE, I: HCD at 30 NCE, J: HCD at 35 NCE, K: HCD at 40 NCE, L: HCD at 48 NCE, M: HCD at 50 NCE, N: sceHCD at 25±15 NCE, O: sceHCD at 30±10 NCE, P: sceHCD at 30±18 NCE, Q: sceHCD at 35±15 NCE. The solid bar on each box plot represents the median and a circle shows the mean. Data are from PXD023448.^45^ **B)** Figures from Maliepaard et al. were recreated using data from GlyCounter. Line plots show the relative intensity of HexNAc (m/z 204.09) and Hex-HexNAc (m/z 366.14) ions across different HCD NCEs in isomer A (far left), which has an antennary galactose on the α1,3-linked mannose, and isomer B (middle left), which has an antennary galactose on the α1,6-linked mannose. The heatmaps show the relative intensities of HexNAc (middle right) and Hex-HexNAc (far right) across different HCD NCEs for mixes of isomers A and B. Data are from MassIVE repository MSV000091172.^47^ **C)** Boxplots show the distribution of percent TIC of HexNAc and Hex-HexNAc ions when glycopeptides are fragmented with ETD, EThcD, and activated ion electron transfer dissociation (AI-ETD). Data are from PXD011533.^49^ **D)** Boxplots show percent TIC for multiple HexNAc-related oxonium ions (HexNAc – CH_6_O_3_, m/z 138.06; HexNAc – C_2_H_4_O_2_, m/z 144.07; and HexNAc, m/z 204.09) and sialic acid-related oxonium ions (NeuAc – H_2_O, m/z 274.09 and NeuAc, m/z 292.1027) when fragmented with HCD or ultraviolet photodissociation (UVPD). Data are from MassIVE database MSV000094544.^50^

Because of the chemical complexity of glycopeptides, glycopeptide fragmentation has received considerable attention. We wanted to show how GlyCounter can aid interpretation of promising alternative fragmentation methods that still require further development for widespread adoption. One such method is activated ion-electron transfer dissociation (AI-ETD), which uses low-energy infrared photons for supplemental activation to improve sequence-information fragment ion yield in ETD reactions.^48^ Laser power can be used in AI-ETD to not only generate peptide sequencing ions through boosted ETD efficiency, but also to produce glycan-specific fragments through cleavage of glycosidic bonds via vibrational activation. This hybrid glycan and peptide fragmentation is similar to EThcD (which uses collisions instead of photons), but they differ in the timing of activation and energy input. We used GlyCounter to re-examine AI-ETD and EThcD spectra^49^, finding that AI-ETD produces more signal for both HexNAc (m/z 204.087) and HexNAcHex (m/z 366.140) oxonium ions (**Figure 2C**). We were also interested to test GlyCounter’s ability to probe spectra from another photoactivation technique, ultraviolet photodissociation (UVPD). Using GlyCounter to explore data from Helms et al.^50^, we saw that HCD generates higher signal in lower m/z oxonium ions HexNAc - CH6O3 (m/z 138.055) and HexNAc - C2H4O2 (m/z 144.066), while UVPD spectra contain much higher levels of HexNAc, which could be useful when designing product-ion dependent data acquisition schemes. Our analyses of these dissociation methods with GlyCounter underscore its value for investigating nuanced fragmentation patterns from multiple MS/MS approaches that hopefully will be helpful as the glycoproteomics field continues to explore this space

### Glyco-enzyme evaluation

Glycoproteomic workflows often employ enzymes called glycosidases to cleave glycosidic bonds between monosaccharides, simplifying the glycan for both analytical and biological purposes. The efficiency of these enzymes is an important, but commonly overlooked, parameter for these experiments, especially as glycoproteomic search engine settings rely on the enzymes producing their desired effect to correctly identify the glycopeptide. PNGaseF, arguably the most popular glycosidase for glycoproteomics (or in this case, de-glycoproteomics), is an endoglycosidase that cleaves the bond between the asparagine and the glycan in N-linked glycopeptides to remove the glycan completely. This cleavage creates deamidation at the asparagine, generating a mass difference of 0.984 Da and simplifying the (de)- glycopeptide for further analysis. PNGaseF reactions can be checked by gel-based assays, but this check is not always included. Instead, conditions that were previously used by other labs or with other lots of enzymes are assumed to be sufficient for new experiments, which may not always be the case. As a proof-of-principle of how GlyCounter can be used to quickly test for enzyme activity, we evaluated PNGaseF-treated fetuin samples from Sun et al.^51^ **Figure 3A** demonstrates this check with a significant decrease in the percentage of total MS/MS scans with oxonium ions following a PNGaseF treatment of fetuin. Oxonium ion signal after enzymatic treatment could be attributed to inefficient PNGaseF activity, but it is likely signal derived from known O-glycosites on fetuin (**Supplementary Figure 10**).^52^ Similar tests can be done with another common enzyme in glycoproteomics, sialidase, which removes terminal sialic acid residues from glycans. We examined the effects of sialidase on O-glycopeptides on multiple highly glycosylated mucin-domain glycoproteins (**Figure 3B**).^53^ GlyCounter shows that the sialic acid-specific oxonium ions effectively disappear from spectra after sialidase treatment; in this study, this was especially important information for testing cleavage motifs of O-glycoproteases. GlyCounter also revealed useful information about sialic acid types present in the sample. At first glance, these are all recombinant human proteins, meaning they would be expected to have mainly Neu5Ac and not Neu5Gc. GlyCounter shows clear signal for Neu5Gc-containing glycans for CD43 and GP1ba. A quick check reveals that these proteins were expressed in an NS0 murine cell line, resulting in the presence of Neu5Gc, instead of the CHO-produced MUC16 and PSGL- 1 that only have Neu5Ac. Knowing that both Neu5Ac and Neu5Gc are present in the data is important for database searching, since most search engines require a glycan database input. Assuming the human proteins only contained human-expressed monosaccharides would limit identifications to exclude any spectra that contain NeuGc; in this case that would result in missing out on 30% more glycopeptide identifications that can contribute to defining O-glycoprotease cleavage motifs. In both PNGaseF and sialidase case studies, GlyCounter served as an easy and fast sanity check to make sure critical experimental conditions and enzymes are working in the intended way.

**Figure 3.**
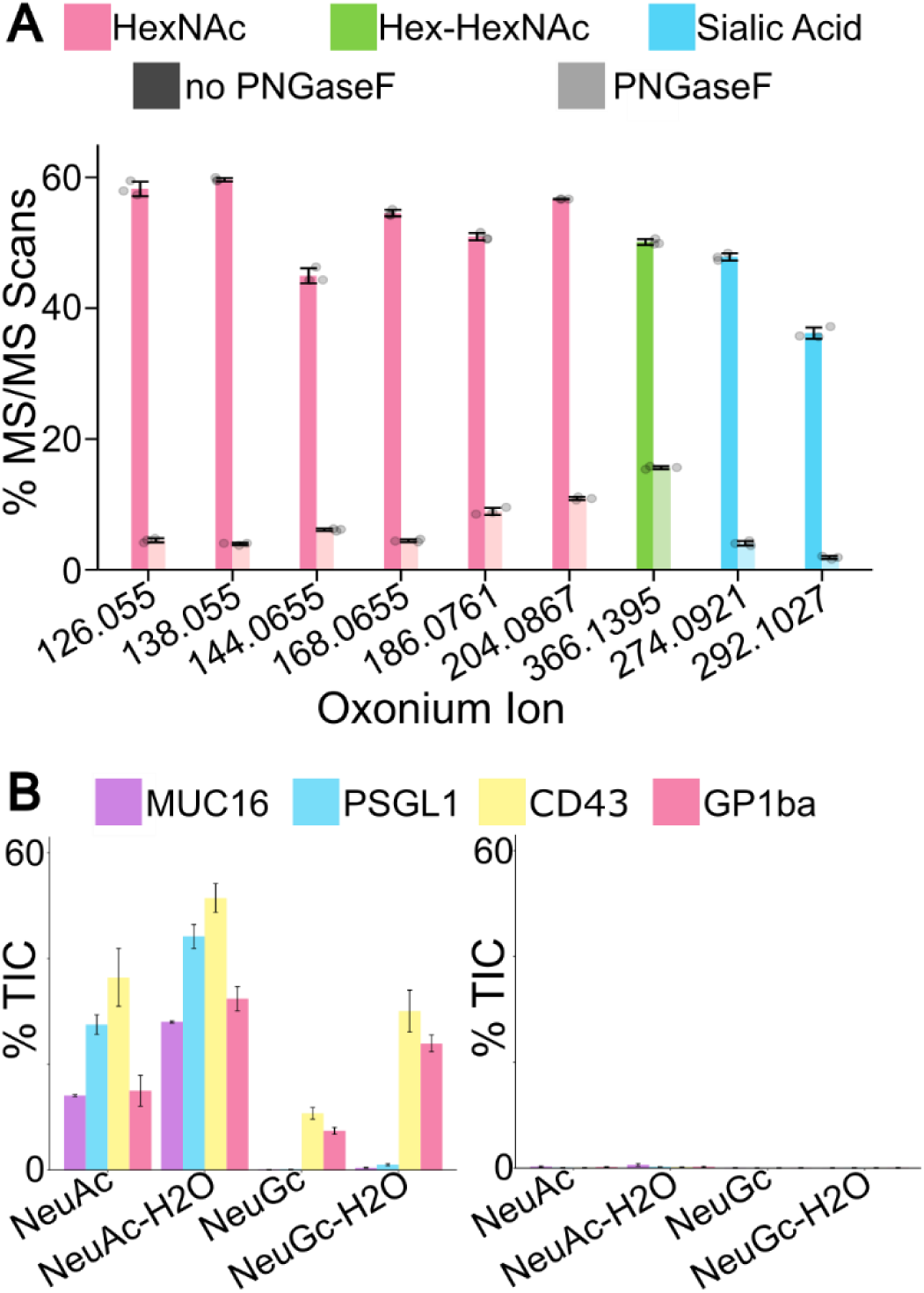
GlyCounter for glyco-enzyme evaluation. **A)** The bar graph illustrates the change in oxonium ion presence before (dark) and after (light) treatment with PNGaseF. Bar height is the average of three runs and the black lines represent one standard deviation. Data are from PXD001571.^51^ **B)** The bar graph shows the average percent TIC of sialic acid ions before (left) and after (right) treatment with sialidase. Bar height is the average of two runs and the black lines represent one standard deviation. MUC16 and PSGL1 were expressed in Chinese hamster ovary cells, while CD43 and GP1ba were expressed in NS0 cells, explaining the presence of NeuGc ions in CD43 and GP1ba spectra. Data are from PXD035775.^53^

### Enrichment efficiency

Glycan heterogeneity makes glycopeptide enrichment a challenge and a critical decision point in glycoproteomics experiments.^54^ We used GlyCounter to examine the oxonium ion content from several enrichment strategies, showing how glycan-specific ions can be just as informative, if not better-suited, than glycopeptide identifications for understanding what enrichment method may be best for a given experiment. First, we analyzed a head-to-head comparison of strong cation exchange (SCX) and high-pH reversed phase offline fractionation methods for phosphoproteomics (**Figure 4A**).^55^ Immobilized metal affinity chromatography (IMAC) used for phosphopeptide enrichment are known to also enrich glycopeptides^56^, but how many glycopeptides are lurking in phosphoproteomics data remains unknown and inconsistent. We were also curious if differences in glycopeptide content could be seen from the different fractionation methods. GlyCounter revealed significant glycopeptide signal in these phosphopeptide-focused datasets, including nearly 14% of all spectra in the optimized high-pH enrichment being classified as “likely glycopeptide”. We also found that SCX less consistently separates IMAC-enriched glycopeptides, with the largest fraction of glycopeptides eluting late in the gradient, likely during the wash. While there are many interesting directions to consider from this simple evaluation, our key take away is that GlyCounter can inform analysts when relatively high numbers of glycopeptide MS/MS spectra are present in unexpected datasets. Conversely, we also examined data from Čaval et al. that used IMAC to enrich phosphorylated glycans containing mannose-6-phosphate (M6P) to study their effects on lysosomal processes.^57^ A key feature of their method is the ability to tune iron IMAC to enrich M6P-glycans and not sialylated glycans, so we used GlyCounter to quantify M6P- and sialic acid-specific ions between their wild-type (WT) cells and cells deficient in Acp2 and Acp5, i.e., acid phosphatases targeting M6P. As reported by Caval et al., we saw a large number of spectra containing Man-P and Man-2P in the KO cell line (**Figure 4B**). Differing from the IMAC study above, and as reported by the authors, few spectra with sialic acid-specific glycans were detected.

**Figure 4.**
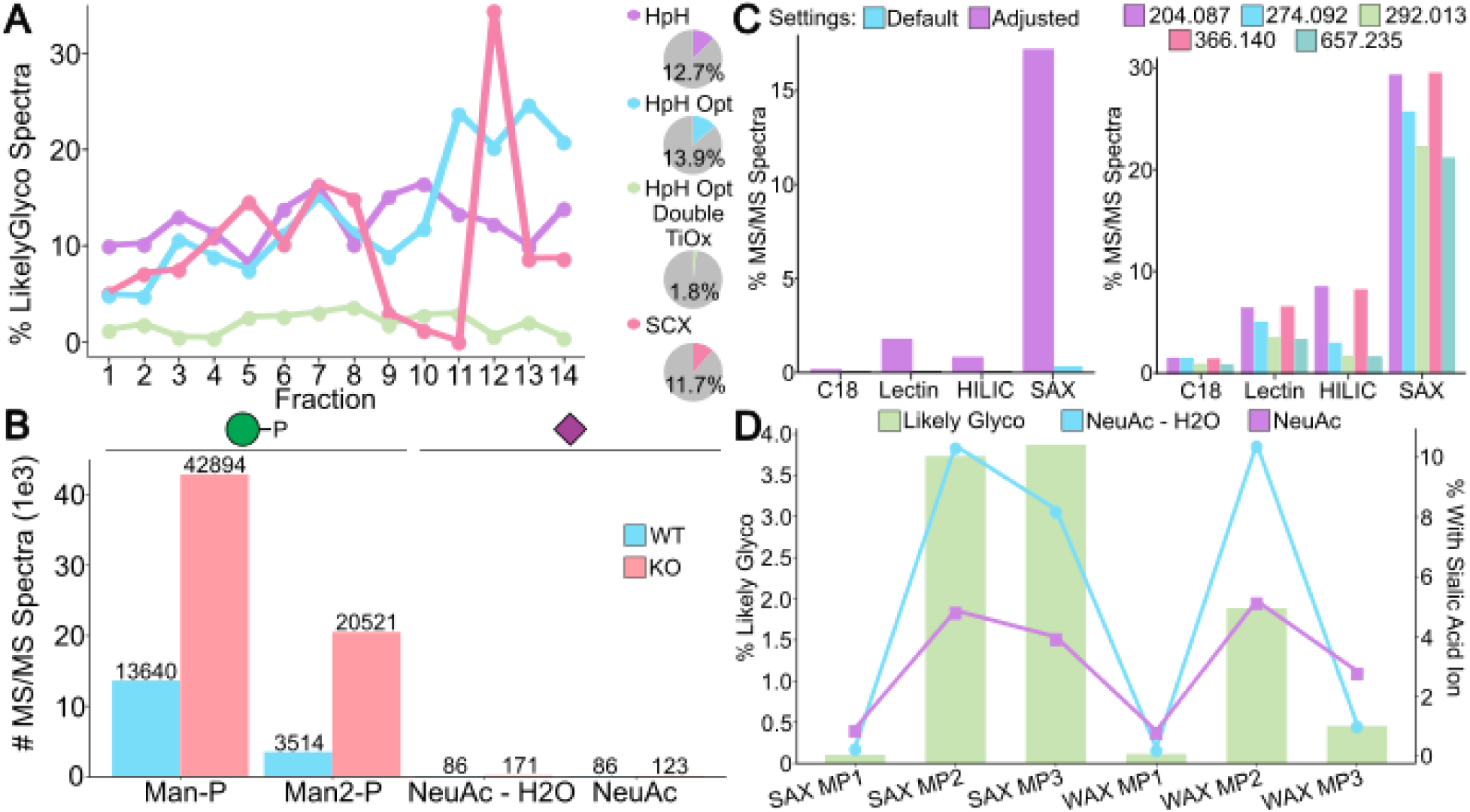
Enrichment techniques evaluated with GlyCounter. **A)** Line graphs display percent likely glycopeptide spectra from GlyCounter for each fraction of four different enrichment/fractionation methods test for phosphoproteomics methods. The chromatographic methods are high-pH (HpH), optimized high-pH (HpH Opt), optimized high-pH with double titanium dioxide (HpH Opt Double TiOx), and strong cation exchange (SCX). Pie charts show the total percent likely glyco across all fractions. Data are from PXD001404.^55^ **B)** The bar graph displays the number of MS/MS spectra that mannose-phosphate and sialic acid ions appear in before and after enriching for mannose-6-phosphate. The wild-type (WT) cells show much lower amounts of mannose-phosphate (Man-P, m/z 243.03 m/z) and mannose-2-phosphate (Man2- P, 405.08 m/z) than the Acp2 and Acp5 knockout (KO) cells. The iron IMAC enrichment method used in this study minimized the number of sialic acid enriched, as shown by few spectra with NeuAc (m/z 292.10 m/z) and NeuAc – H_2_O (m/z 274.09). Data are from PXD010333.^57^ **C)** Bar graphs compare oxonium ion data for four enrichment methods. The enrichment methods are C18 reverse phase, multiple lectin affinity chromatography, hydrophilic interaction liquid chromatography (HILIC), and strong anion exchange- electrostatic repulsion hydrophilic interaction chromatography (SAX). The graph on the left shows percent likely glycopeptide spectra from GlyCounter for each of the enrichment methods with both default settings that assume a lower m/z range below m/z 126 (blue) and results using adjusted settings that only require four oxonium ions in the top 25 peaks instead of the default eight (purple). SAX shows the highest percent likely glyco out of the four methods. Right: The graph shows the percent of MS/MS scans that certain oxonium ions are detected in for each enrichment method. SAX also shows the highest percentage of MS/MS scans where individual oxonium ions occur. Data are from PXD005655.^58^ **D)** The combined graph shows the percentage of likely glycopeptide spectra from GlyCounter (bar) for three mobile phase buffers for both strong anion exchange (SAX) and weak anion exchange (WAX). The line graphs show the percentage of MS/MS spectra with specific sialic acid ions, NeuAc and NeuAc – H_2_O. Data are from PXD022988.^59^

We next looked at an N-glycopeptide enrichment comparison study from Totten et al.^58^ Despite seeing many oxonium ions from different enrichment methods, GlyCounter’s default settings classify very few spectra as “likely glycopeptide” (**Figure 4C**) due to the low m/z scan range boundary (m/z 190) precluding detection of several common oxonium ions (e.g., m/z 138, 168, 186). The flexibility of GlyCounter lets us tailor the analysis to the experiment and quickly re-run the analysis with adjusted settings requiring only 4 oxonium ions instead of the default 8. We examined what effect raising the lower m/z bound would have and found that the results are comparable to a full scan range when using these adjusted settings (**Supplementary Figure 11)**. Ultimately, this flexibility allowed us to get a clearer picture of the results, and our analysis supports the original conclusions that strong anion exchange electrostatic repulsion hydrophilic interaction (SAX-ERLIC) solid-phase extraction is a favorable enrichment method for N-glycopeptides in plasma. Finally, we further investigated enrichment with ERLIC using data from Cui et al. comparing SAX-ERLIC to weak anion exchange (WAX)-ERLIC.^59^ **Figure 4D**, indicates that while SAX is more effective at broadly enriching glycopeptide species (more oxonium ions, hence more likely glycopeptide spectra), the right buffer conditions for WAX can enrich sialylated glycopeptides with a similar efficiency as SAX, likely due to the negative charge on sialic acids. GlyCounter also revealed that while effective at improving glycoproteome sampling, these ERLIC methods are not necessarily efficient, as shown by a small fraction of total MS/MS scans being assigned as likely glycopeptide. In all cases, we arrived at conclusions consistent with the full study with only minimal analysis time and without having to search, identify, or further process glycopeptide data.

### GlyCounter as a supplement to database searching

While some glycoproteomics-centric search engines can leverage an open-modification search or glycan-database free search^35,39,60,61^, most algorithms use a glycan database to define the possible glycan compositions to consider. Approaches to assign peptide and glycan moieties differ in which species they prioritize first, and heterogeneity in strategies leads to considerable variability in glycopeptide identifications between search engines.^8^ Our group has found that supplementing search algorithm identifications with concrete spectral data has helped evaluate various search approaches. To demonstrate, we first compared data from popular search algorithms against GlyCounter’s default likely glycopeptide spectrum output (**Figure 5A**).^35,37,62,63^ We observe a much larger number of spectra considered likely to be a glycopeptide by GlyCounter than identifications by the database search algorithms. Since GlyCounter does not take peptide fragments into account for its “LikelyGlyco” characterization, it is probable that many of these spectra did not have sufficient peptide information for a search algorithm to make an identification. The high abundance of B- and Y- type ions often means that peptide b- and y-ions are much less intense than in a non-modified peptide spectrum, which leads to lower scores that may not pass FDR control in so-called “peptide-first” database searching. GlyCounter shows how many spectra we are potentially missing in our database searches that could be glycopeptide species. An additional layer of complexity when including glycan databases is whether to consider covalent glycan modifications like acetylation, sulfation, or metal ion adducts. We used GlyCounter to evaluate the presence of oxonium ions that can indicate the presence of modified glycans in a recently published study from Liu et al. that investigated acetylated sialic acids from mouse and rat sera compared to human serum (**Figure 5B**).^64^ Our analysis shows that their conclusions about sialic acid acetylation from StrucGP glycopeptide identifications can be fully recapitulated from only considering diagnostic oxonium ions for nonmodified and acetylated sialic acids. Therefore, using GlyCounter to inspect raw data for these ions can add value to knowing whether to include glycan modifications in glycopeptide searches and as useful features to validate glycopeptide spectral matches.

**Figure 5.**
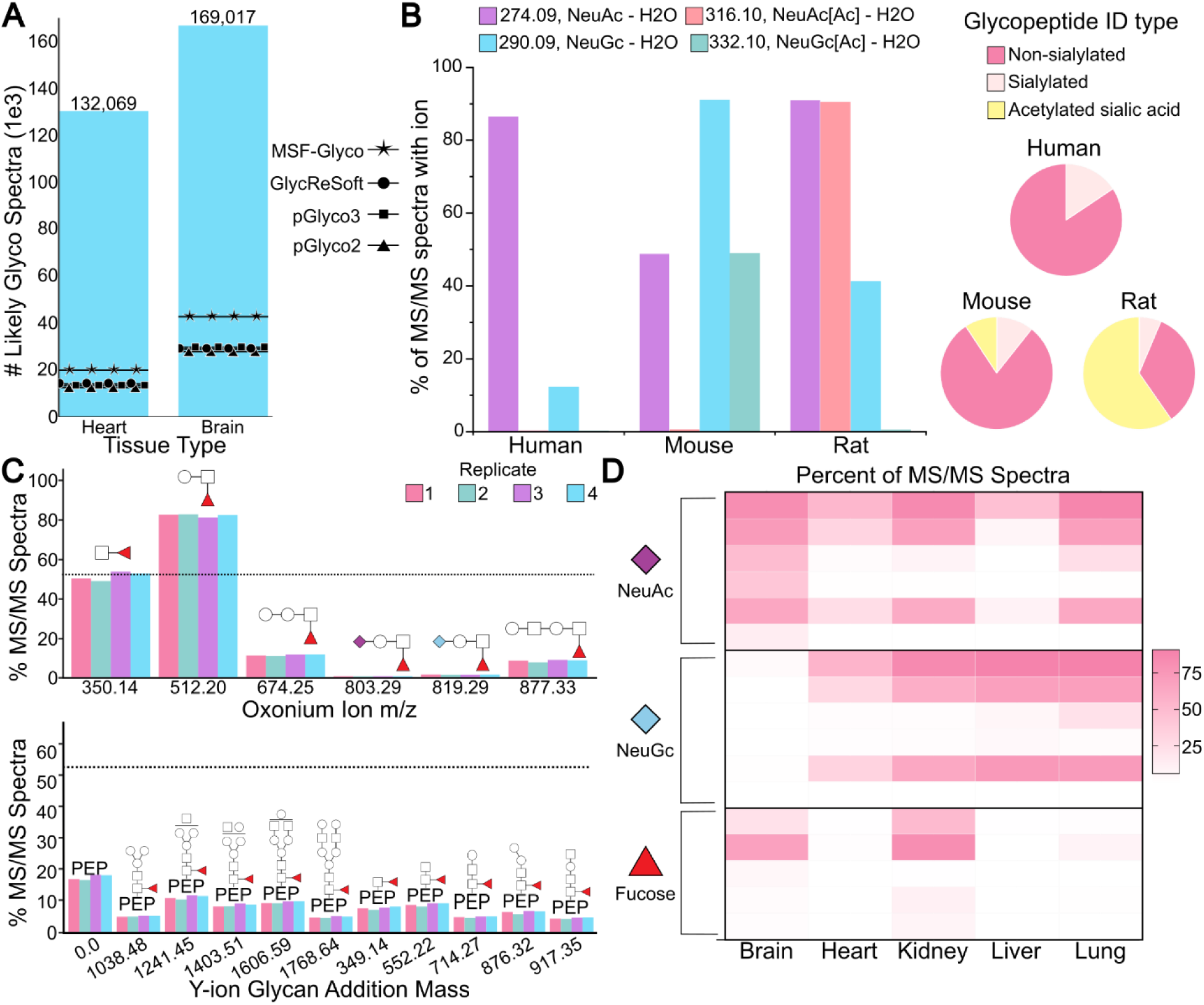
GlyCounter to supplement database searching. **A)** The same mouse tissue glycoproteome dataset was searched with four different search engines and GlyCounter to compare the number of glycopeptide spectral matches (search engines) and number of likely glycopeptide spectra (GlyCounter). The search engines compared are MSFragger-Glyco (MSF-Glyco)^35^, GlycReSoft^63^, pGlyco3^37^, and pGlyco2^62^. Data are from PXD005411 (brain) and PXD005413 (heart).^62^ Search engine results are taken from previously published/reported data. **B)** Pie charts show the percentage of identifications from human, mouse, and rat serum with three different N-glycopeptide types: not sialylated, sialylated, and containing acetylated sialic acid. We also plot a bar graph, which shows the percentage of MS/MS scans containing the NeuAc – H_2_O (m/z 274), NeuAc[Ac] – H2O (m/z 316), NeuGc – H2O (m/z 290), and NeuGc[Ac] – H2O (m/z 332) oxonium ions, which are all pre-loaded in GlyCounter. Data are from PXD053293^64^ and include search results from StrucGP. **C)** The bar graph shows the percent of MS/MS spectra with fucose-specific oxonium and Y-type ions detected with GlyCounter and Ynaught in mouse kidney tissue. Y-ion masses represent the addition of the depicted glycan mass to the unglycosylated peptide backbone. The dotted line at 52.5% represents the percent of spectra that were identified to be fucosylated by pGlyco2. Replicates 1, 2, 4 (here represented as 3), and 5 (here represented as 4) were used from the original data obtained from PXD005412.^62^ **D)** The heatmap displays the percent of MS/MS spectra containing each ion type in five different mouse tissues. The oxonium ions searched for from top to bottom are NeuAc – H_2_O (m/z 274.09), NeuAc (m/z 292.10), NeuAc-Hex (m/z 454.16), NeuAc-HexNAc (m/z 495.18), NeuAc-Hex-HexNAc (m/z 657.23), NeuAc2-Hex-HexNAc (m/z 948.33), NeuGc – H_2_O (m/z 290.09), NeuGc (m/z 308.10), NeuGc-Hex (m/z 470.15), NeuGc-HexNAc (m/z 511.18), NeuGc-Hex-HexNAc (m/z 673.23), NeuGc2-Hex-HexNAc (m/z 964.33), HexNAc-dHex (m/z 350.14), Hex-HexNAc-dHex (m/z 512.20), NeuAc-Hex-HexNAc-dHex (m/z 803.29), Hex2-HexNAc-dHex (m/z 674.25), and Hex2-HexNAc2-dHex (m/z 877.33). Data are from PXD005411 (brain), PXD005413 (heart), PXD005412 (kidney), PXD005553 (liver), and PXD005555 (lung).^62^

We were also curious if GlyCounter results could show trends in “glycotyping” tissues in large datasets that previously required significant search algorithm development. In the paper describing pGlyco2, the authors found that 52.5% of spectra from mouse kidney tissue contained fucosylated glycopeptides,^62^ so we used GlyCounter and the Ynaught module within GlyCounter to search for oxonium and Y-type ions that contain fucose. This case study showcases the Ynaught module, which pairs with glycopeptide identifications to easily extract Y-type ions from MS/MS spectra, allowing us to examine multiple glycopeptide features beyond only sugar fragments. **Figure 5C** compares ion signal extracted by GlyCounter (bars) with the percentage of fucose-containing glycopeptide identifications in kidney from pGlyco2 (line). The presence of a variety of ions confirms a relatively high degree of fucosylation in the tissue, although the HexNAc- dHex (m/z 350.14) is the only ion that faithfully recapitulates the percentage seen with glycopeptide identification. Interestingly, Hex-HexNAc-dHex (m/z 512.20) appears in over 80% of spectra collected from the mouse kidney tissue, indicating that some “diagnostic” ions must be carefully evaluated because they can be generated from numerous epitopes. A second interesting trend from that study showed Neu5Ac-containing glycopeptide ions detected in every tissue, while Neu5Gc-containing glycopeptides were noticeably absent in brain tissue. **Figure 5D** plots the percentage of MS/MS spectra that contain sialylated and fucosylated oxonium ions from all five tissues analyzed in the study. These data confirm trends from the glycopeptide identifications, including fucose ions detected mostly in brain and kidney tissues and Neu5Gc ions absent from brain tissue. GlyCounter data allows us to rapidly glycotype these tissues, providing a framework for validating glycopeptide identifications and investigating biological implications of these trends regardless of search engines used.

### Investigating undesired glycopeptide fragmentation during method development

High- field asymmetric waveform ion mobility spectrometry (FAIMS) has emerged as a technique useful in proteomics methods for rapid gas-phase fractionation prior to MS analysis and has even been postulated as a complement, if not even an alternative, to liquid chromatography (LC).^65–67^ Since FAIMS separations are rapid and comparatively sensitive, complex proteomics samples uniquely benefit from additional fractionation and approach comparable depth to that of analogous LC approaches.^68^ These studies frame FAIMS as an attractive strategy for glycopeptide analysis, as glycoforms readily co-elute within a given LC peak. Initial work investigating the complementary nature of FAIMS to other condensed-phase separations reported a 25% increase in a glycoproteome with a primary focus on N-glycopeptides.^69^ Rangel-Angarita et al. examined the use of FAIMS for O-glycopeptide identification^70^. While their FAIMS experiments identified 2- to 5-fold increases in spectral matches, further investigation showed that in-FAIMS fragmentation was a major contributor to the increased identifications. In the simplest of the analyzed samples, digested podocalyxin, 24.1% of the glycopeptide spectral matches resulted from in-FAIMS fragmentation (IFF). As IFF is challenging to identify without a robust manual investigation into individual spectra and their accompanying chromatograms, we wanted to investigate if oxonium signal alone from GlyCounter would suggest similar conclusions.

After searching the SmE-digested podocalyxin data with GlyCounter, we found similar trends between likely glycopeptide spectra, glycopeptide spectral matches (glycoPSMs), and unique glycopeptides as seen in **Figure 6A**. Interestingly, we see almost twice as many likely glycopeptide spectra in a run without FAIMS than at the most effective FAIMS compensation voltage (CV). In **Figure 6B**, the 17 most abundant oxonium ions are shown with their presence in HCD scans normalized to the HexNAc ion at each CV. Even after normalizing to estimate the presence of glycopeptides, there is some variability in other oxonium ions. To examine this further, **Figure 6C** shows the trends for sialic acid ions and Hex-HexNAc relative to the presence of HexNAc and glycoPSMs. The Hex-HexNAc ion clearly follows the trend of the glycoPSMs whereas the two sialic acid ions deviate. The NeuAc-H_2_O ion frequency peaks at -45 V and rapidly decreases at higher CVs. NeuGc on the other hand, remains closest to the HexNAc signal. In general, data from GlyCounter aligns with the conclusions drawn by the publication that FAIMS may complicate O-glycopeptide identifications by artificially increasing microheterogeneity by IFF.

**Figure 6.**
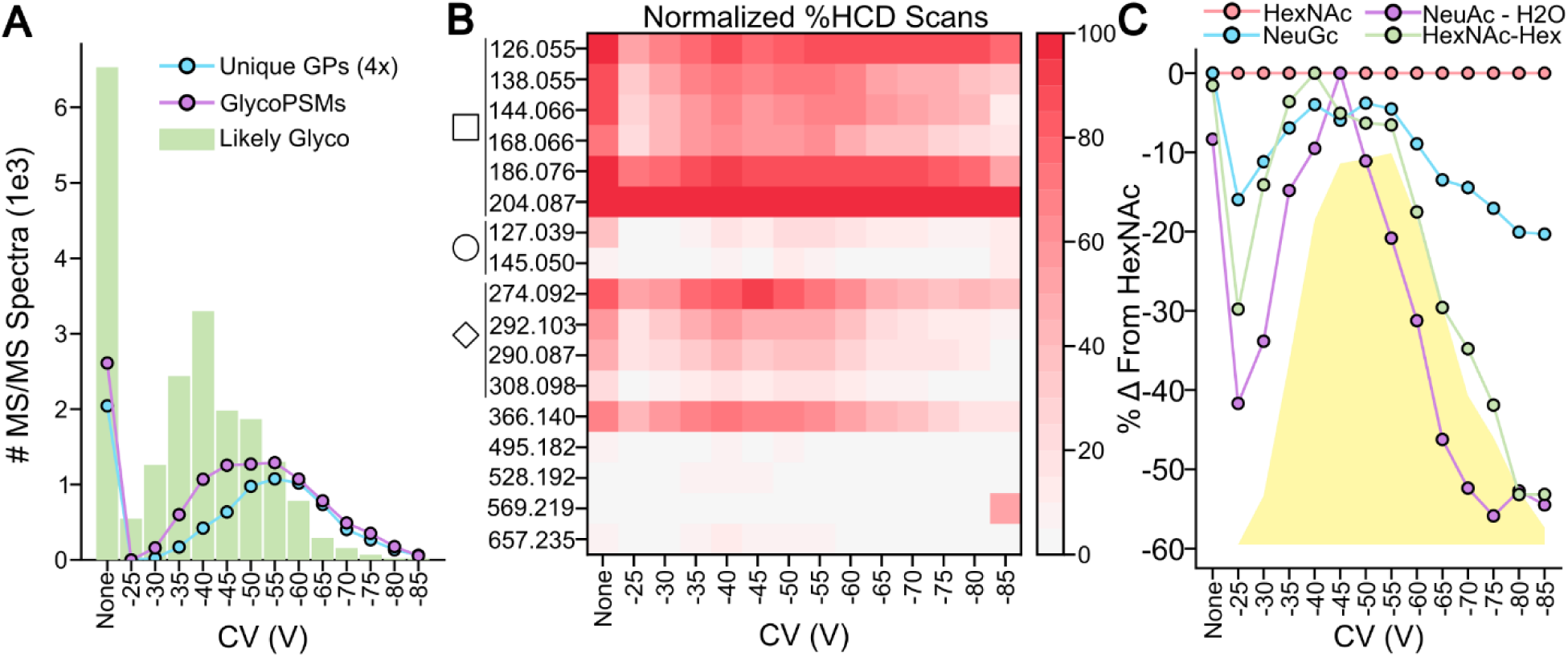
Using GlyCounter to evaluate FAIMS for glycopeptides. **A)** Number of MS/MS spectra that were assigned likely glycopeptide by GlyCounter for many compensation voltages (CV) in a FAIMS method. “None” means that no FAIMS was used. The number of glycopeptide spectral matches (glycoPSMs) and unique glycopeptide identifications (unique GPs) are overlayed to show that the trends between them line up. The number of unique GPs was scaled up by a factor of 4. **B)** The heatmap shows the individual ions that are present in HCD scans at different CVs. The data is normalized to HexNAc (204.09 m/z) being present in 100% of spectra. Oxonium ions have been categorized as being derived from HexNAc (square), Hex (circle), sialic acid (diamond), or general oligosaccharides (366.140, Hex-HexNAc; 495.182, NeuAc- HexNAc; 528.192, Hex2-HexNAc; 569.219, Hex-HexNAc2; 657.235, NeuAc-Hex-HexNAc). **C)** The line graphs show the difference in normalized percent of HCD scans between HexNAc and three ions of interest. The yellow trace represents the number of glycoPSMs found per CV. We see that Hex-HexNAc (m/z 366.14 m/z) follows the trend of the glycoPSMs, while the two sialic acid ions (NeuAc – H_2_O, m/z 274.09 and NeuGc, m/z 308.10) do not. Data are from PXD041217.^70^

### Y-type and Custom Ions

One key feature of GlyCounter is its ability to accept custom ion inputs to tailor analyses to specific experiments. We first wanted to use this feature to investigate glycan formylation, a phenomenon described by Zhi et al., where glycans gain a +28Da modification when glycopeptides are stored in formic acid.^71^ We created modified oxonium ions representing formylated glycans/glycan fragments as a custom input (**Supplementary File 3**) and used GlyCounter to examine their prevalence. GlyCounter results in **Figure 7A** show that after 14 days stored at -20°C, as much as 14.5% of the oxonium ion total ion current consists of formylated oxonium ions. Here, GlyCounter can serve as a useful screening tool to track this undesired modification and reduce errors in sample preparation. A second, common use case for custom oxonium ions is glycan labeling via biorthogonal and click chemistry, so we evaluated GlyCounter’s performance for extracting click chemistry-specific oxonium ions in the IsoTaG- labeled O-GlcNAc dataset from Woo et al.’s study on human T cells.^72^ In addition to a few common oxonium ions, this tag creates a “HexNAzoSi” oxonium ion at m/z 345.14, which we uploaded to GlyCounter as a custom ion (**Supplementary File 3**). In that study, cells were cultured under three stimulation conditions and a separate control was treated with DMSO. **Figure 7B** shows the percent of spectra containing HexNAzoSi ion from the unlabeled control and each of three different conditions. Effectively zero MS/MS spectra in the unlabeled sample contained this ion (likely only signal that can be attributed to noise). The IsoTag labeling approach appears to have successfully worked in the three treatment conditions, as each shows at least 9% of spectra containing the HexNAzoSi ion. GlyCounter’s custom ion function is easily adapted to individual experiments, opening potential use-cases beyond standard oxonium ions.

**Figure 7.**
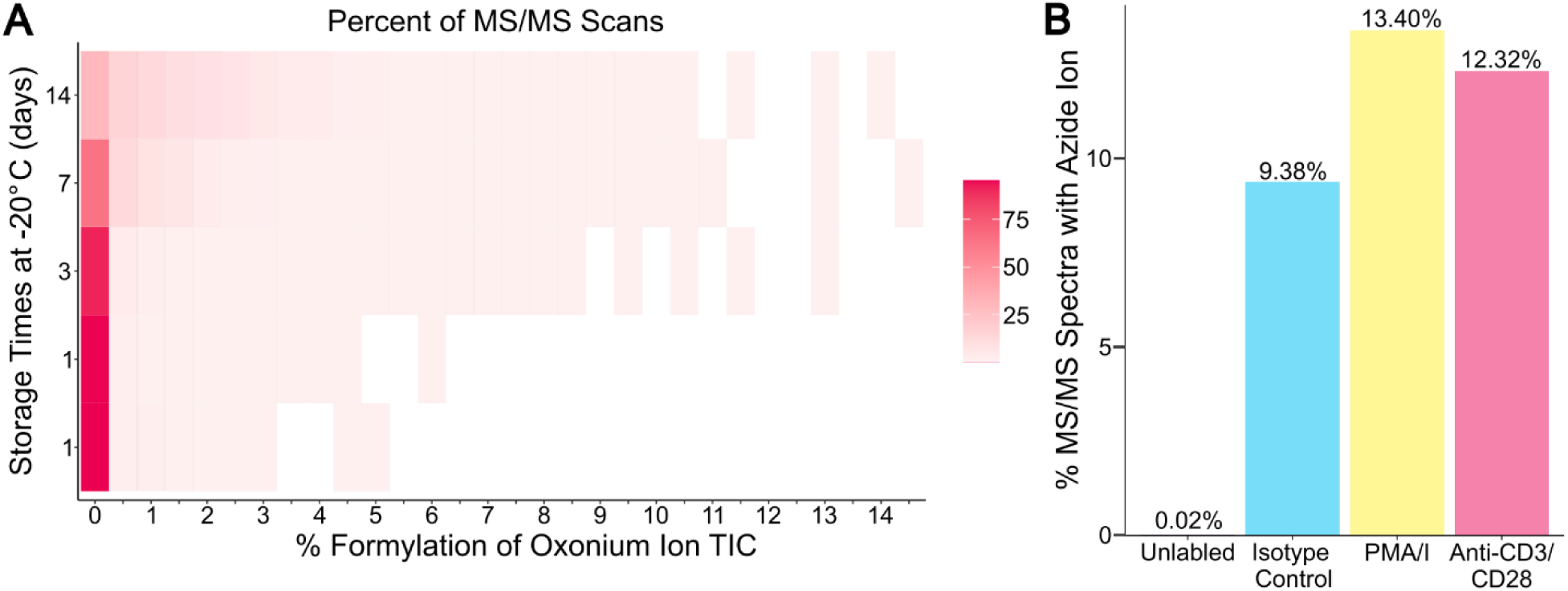
Extracting modified glycan ions using the custom ion upload in GlyCounter. **A)** The heatmap shows the binned percent of oxonium ion TIC coming from formylated oxonium ions (x-axis) and the percent of MS/MS scans per bin (heat) for each amount of time (y-axis) that the glycopeptide sample was stored at -20°C. There were two trials performed at 1 day, then one each at 3, 7, and 14 days. We observed higher percentages of formylated ions starting at 3 days and increasing at the 7- and 14-day timepoints. Data are from PXD023448.^71^ **B)** The bar graph shows the percent of MS/MS spectra with the IsoTaG derived azide oxonium ion (345.14 m/z) between the four experimental conditions. The unlabeled control sample with no tag should not contain this ion, and the other sample types contain the tag at various levels. Data are from PXD004559.^72^

## DISCUSSION

Glycan-specific ions, especially oxonium ions, are commonly used for several purposes in glycoproteomics and beyond, including but not limited to: determining the presence of glycopeptides in a dataset, examining gradient suitability for glycopeptide separations, classifying glycans based on known fragmentation ratios, evaluating glycopeptide identifications, and potentially informing glycan structure. Most studies rely on manual inspection to understand the oxonium ion content in their data, or they must use glycopeptide identifications to inform their inspection of oxonium ions. We developed GlyCounter as an automated tool to enable straightforward access to more than 50 oxonium ion features to make exploring glycopeptide content in raw mass spectra more accessible. We now use GlyCounter in our group for many method development and data analysis efforts, and because it has been so useful, we want to bring this user-friendly tool to the broader glycoproteomics community. To highlight how GlyCounter might be useful, we highlighted several use cases, beginning with examining glycopeptide fragmentation trends using established and emerging dissociation methods. We also showed how GlyCounter can provide rapid access to glycan-specific ion content to screen enzyme efficiency, evaluate glycoproteomic enrichment methods (and methods where glycopeptides may be unknowingly or unintentionally co-enriched), and explore features in complex glycoproteomics datasets that can either improve database search strategies or validate search results. Instrumentation and method development is still an active field, so we also wanted to highlight how GlyCounter can help understand strengths and weaknesses of new tools, e.g., FAIMS. Finally, we know that the glycoproteomics community looks at diverse sample types with heterogeneous glycan features and experimental designs. Given this, we wanted to make GlyCounter as flexible and useful as possible, so we enabled user-defined custom ion uploads and showcased this utility for modified glycan case studies. We also added a Ynaught module that uses glycopeptide identifications as input to look for Y-type ions – which are often difficult to extract with current software for further post-search processing. Additionally, various experiment designs and instrumentation will have different analytical requirements, so we gave users the option to choose filtering and evaluation metrics that best suit their needs in case our default settings are not suitable. Although we did not provide examples of data independent acquisition here, GlyCounter is compatible with these workflows. Note, for standard proteomic experiments that use MS/MS scan ranges that start at higher m/z values, users will likely need to adjust parameters accordingly to look for oxonium ions with appropriate m/z values and to change scan settings for LikelyGlyco evaluations. That said, we do not expect standard proteomics data to contain an abundance of glycopeptide data because enrichment is typically required to deal with signal suppression issues.^54^

All evaluations and case studies in this manuscript are from published datasets, highlighting 1) the flexibility of GlyCounter and 2) the value of the glycoproteomics community making their data publicly available. The complexity of glycosylation merits constant and communal data (re)- evaluation, and we appreciate the authors who agree by depositing their data in publicly accessible repository. Importantly, many challenges inherent to glycoproteomic experiments remain and must be considered when using GlyCounter to evaluate data. These include possible contribution of most glycopeptide signal from a few abundant glycoforms (especially in biofluid analysis) and the presence of multiple MS/MS spectra that represent the same glycoform but come from distinct chemical species arising from non-specific protease activity, chemical adducts, and multiple charge states. Thus, interpreting the implications of GlyCounter data and “Likely Glyco” assignments must always be carefully considered in the context of the sample as to not over-interpret results.

In all, our goal is to make glycoproteomics data analysis more straightforward, robust, and accessible to the growing population of glycoproteomics researchers. GlyCounter accomplishes this goal by providing flexible access to explore individual ions and relationships of ions across experimental conditions. Although non-glycosylated sample types are outside of the scope of what we designed this tool to explore, our flexible interface makes it possible to explore non-glycan applications, e.g., looking for phosphorylation losses from precursor ions in phosphoproteomic datasets. We hope GlyCounter can emerge as a straightforward, useful tool to evaluate the glycan content of glycoproteomic data that is decoupled from glycopeptide identification, ultimately enabling refinement in sample preparation, data acquisition, and post-acquisition identifications. It may even prove useful in integrating with newly developed approaches for released glycan and glycomics workflows.^73–76^

## DATA AVAILABILITY

The data analyzed in this study are available at ProteomeXchange Consortium dataset identifiers PXD023448, PXD011533, PXD001571, PXD035775, PXD001404, PXD005655, PXD010333, PXD022988, PXD005411, PXD005413, PXD005412, PXD005553, PXD005555, PXD041217, PXD023448, PXD004559 and MassIVE repositories MSV000091172, MSV000094544, MSV000083070. GlyCounter can be downloaded from: https://github.com/riley-research/GlyCounter/Releases.

## Supporting information

FileS3_CustomIonUpload

FileS4_DataForGraphs

FileS1_ExampleOutputFiles

FileS2_GlyCounterGuide

## ACKNOWLEDGEMENTS

Research reported in this publication was supported by the National Institutes of Health under Award Number R00GM147304 (N.M.R.), by an ASMS Research Award (N.M.R.), by a Royalty Research Fund (RRF) Award (A207842), by a Searle Scholar Fellowship from the Kinship Foundation (N.M.R.), by the National Institute of General Medical Sciences of the National Institutes of Health under Award Number T32GM153507 (K.K.), and by a Washington Research Foundation Postdoctoral Fellowship (E.S). ^¥^Current affiliation: Hannover Medical School, Hannover, Germany. N.M.R. receives support from Thermo Fisher Scientific under a nondisclosure agreement and is a consultant for Augment Biologics.

## AUTHOR CONTRIBUTIONS

K.K.: Data curation; Formal analysis; Investigation; Methodology; Software; Validation; Visualization; Roles/Writing - original draft; and Writing - review & editing.

H.M.S.: Formal analysis; Visualization;

J.H.R.: Formal analysis; Visualization;

E.S.: Roles/Writing - original draft

T.S.V.: Roles/Writing - original draft

R.Z.: Formal analysis; Visualization;

A.G.B.: Formal analysis; Visualization;

V.R.T.: Formal analysis; Visualization; Software

L.E.M.: Formal analysis; Visualization;

L.S.D.: Formal analysis; Visualization;

N.M.R.: Conceptualization; Data curation; Formal analysis; Funding acquisition; Investigation; Methodology; Project administration; Resources; Software; Supervision; Validation; Visualization; Writing - review & editing.

