## Supplementary material for "Extracting informative glycan-specific ions from glycopeptide MS/MS spectra with GlyCounter": FileS2_GlyCounterGuide

### GlyCounter Guide and Tutorial

#### Introduction

This guide provides a walkthrough for using GlyCounter to process mass spectrometry data. For more information about technical details of GlyCounter, see our [GitHub repository](GitHub repository).

#### MSConvert

GlyCounter is compatible with most forms of mass spectrometry data. Thermo .RAW files need no extra processing unless they are collected in profile mode. All other files should be converted to MzML format using [MSConvert](MSConvert) using the settings shown below:

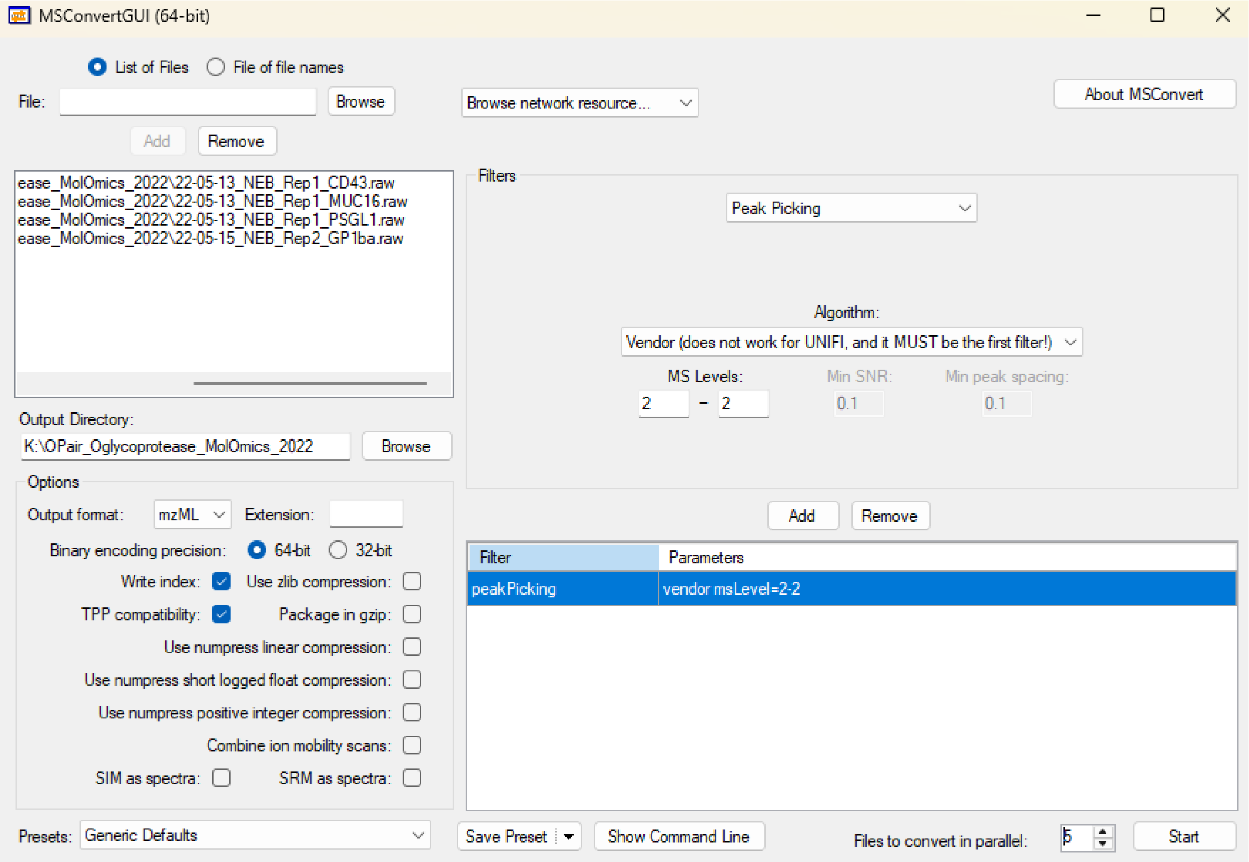


Peak picking is only required if the MS/MS scans are collected in profile mode.

#### Determining Oxonium Ion Selection

Which oxonium ions you choose to look for using GlyCounter is very experiment dependent. In this tutorial, the data chosen is from Figure 3B in the GlyCounter manuscript where samples were analyzed before and after sialidase treatment. In the paper, we chose to look at oxonium ions specific to the sialic acids that would get cleaved by the enzyme to see if the enzyme is working as expected. However, we could also choose to examine all oxonium ions if we were interested in the effect that sialidase has on the entire glycan. If your experiment involves a tag or linker that could create ions that we haven't included in GlyCounter, those ions can be searched for using our custom ions feature. There is an example of the custom ion input spreadsheet in the GitHub repository.

#### GlyCounter User Interface

Step 1: Upload .RAW or .MzML files. This can be achieved by pressing the browse button next to the file upload box (1) or dragging and dropping files into the UI. The text in the box should change to show how many files have been successfully uploaded.

Step 2: Select output directory by pressing the browse button next to the output directory field (2). By default, output files will be placed in the same location as the MS data files that were uploaded.

Step 3: Select relevant ions. The "Check All Ions" (3) and "Check Common Ions" (4) buttons can be used to select many ions at once. Oxonium ions are separated into categories for easier selection of certain glycan types. Custom ion .csv files can be uploaded using the browse button at the bottom of the application (5). To view some suggested negative mode ions, select the "Toggle Negative Mode" checkbox (6).

Step 4: Fragment settings. Fragment ion tolerance can be adjusted in either ppm (default) or Dalton (by clicking the check box labeled Da) (7). The signal-to-noise (S/N) requirement for each peak can also be adjusted, as well as an intensity threshold (8). If noise data is available in the file (Thermo Orbitrap files) S/N will be used. Otherwise, the program will use the intensity threshold.

Step 5: LikelyGlycoSpectrum settings (9). For more information on LikelyGlyco settings, see the manuscript or the GitHub repository. Adjust the settings to your preferences for HCD, ETD (including EThcD), and UVPD scans.

Step 6: MS level settings. Use the arrows to select the range of MS levels that should be searched (10). Note that MS1 searching is possible but would require a custom ion input of the target m/z values to look for.

Step 7: Output IPSA annotations (11). Provides an additional output with ion, m/z, and mass difference columns. This is useful for spectral annotation.

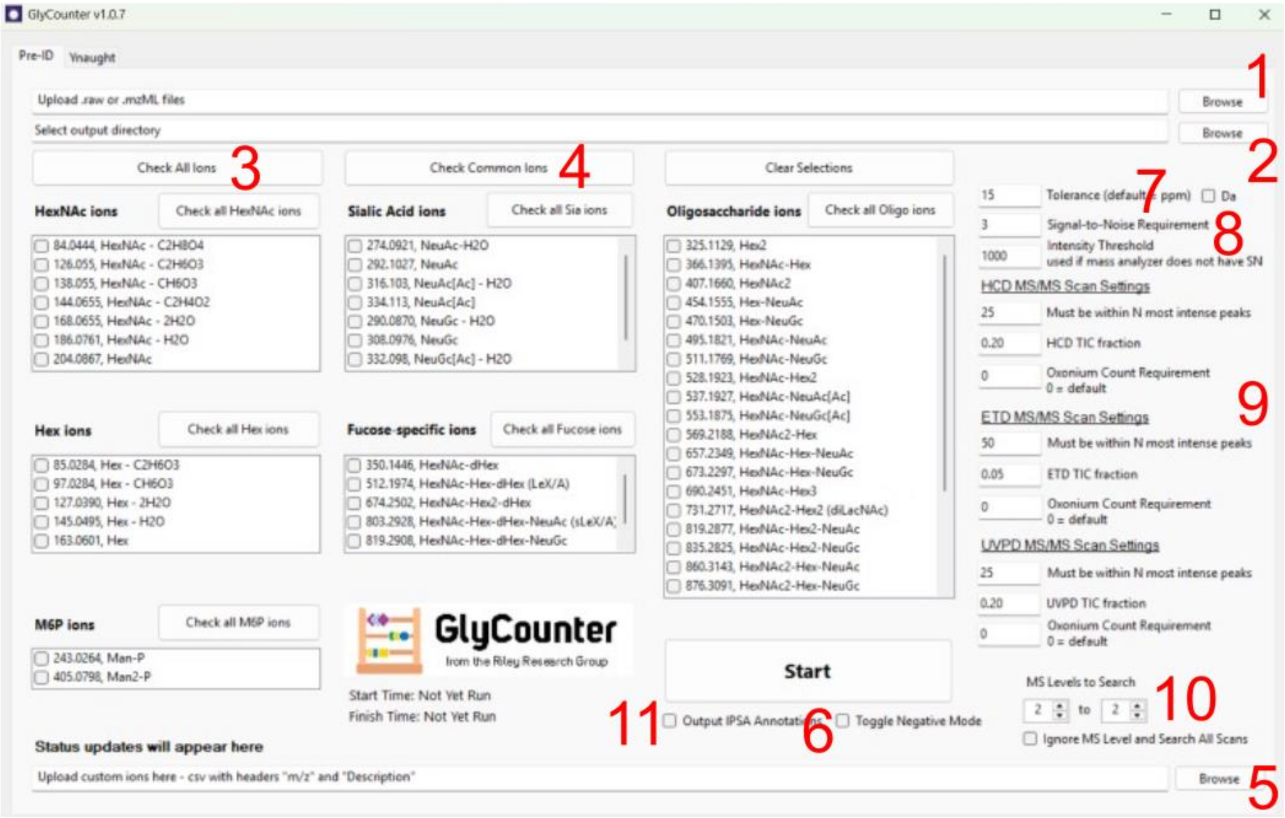


#### GlyCounter Outputs

The GlyCounter outputs contain a lot of data. Here I will show examples of certain types of data to extract from the output files.

**OxoPeakDepth:** The most common use for the peak depth file is to extract peak depths of certain oxonium ions. You can think of peak depth as a ranking of peaks in the spectrum, where 1 is the most abundant peak. In this file, each oxonium ion is given a column so I can examine the distribution of peak depths for each of the sialic acid ions that I'm interested in (See histograms below).

Relating to the peak depth, a summary column towards the end of this file ("OxoInPeakDepthThresh") will show you the number of oxonium ions that showed up in the threshold that was set in Step 5 of the GlyCounter User Interface section of this tutorial. For example, for HCD scans the default setting is to look for oxonium ions in the top 25 peaks. The "OxoRequired" column will then show the minimum number of ions in that threshold required for the spectrum to be considered likely glycosylated.

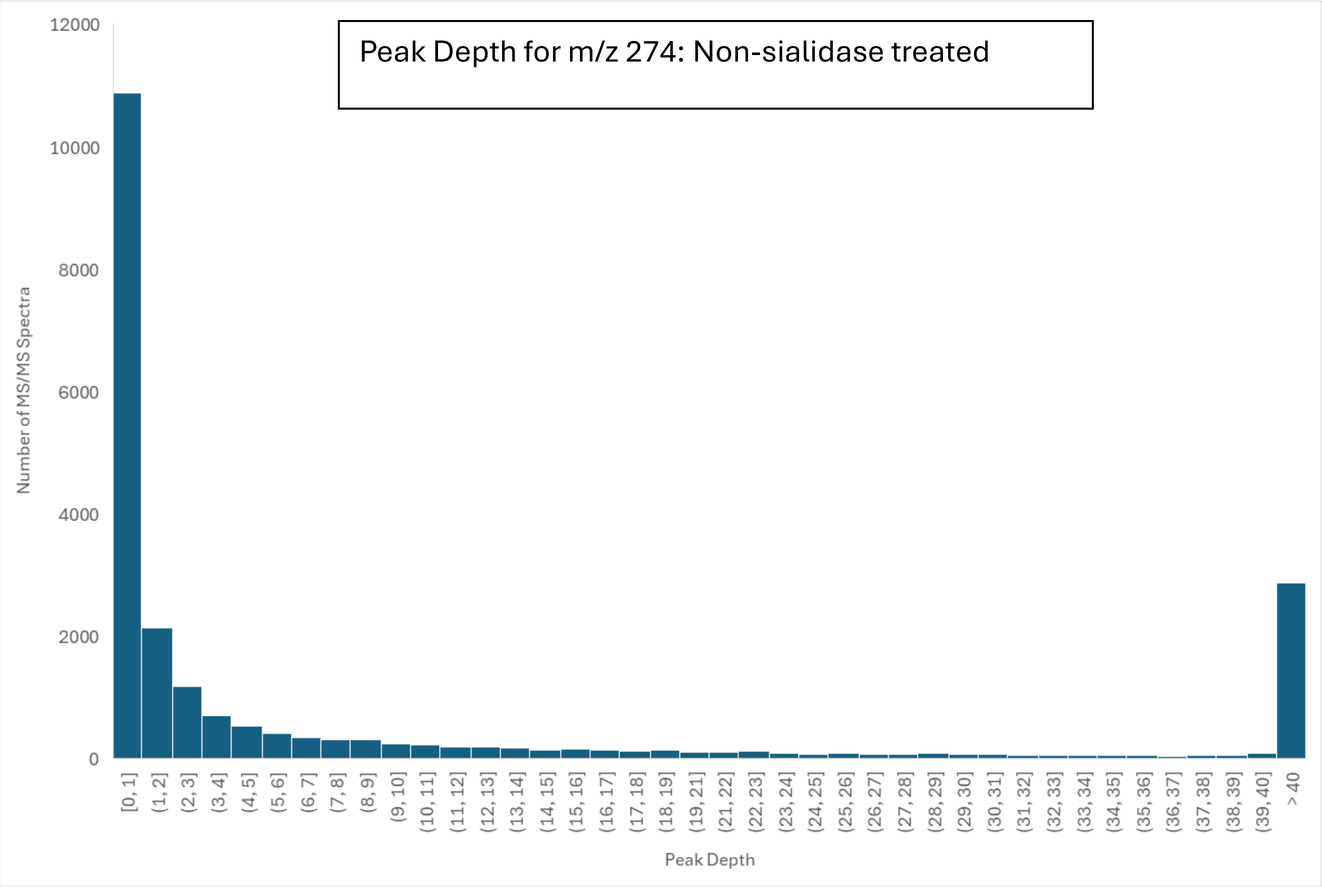
Peak Depth for m/z 274: Non-sialidase treated
Number of MS/MS Spectra
Peak Depth
0
2000
4000
6000
8000
10000
12000
[0, 1]
(1, 2]
(2, 3]
(3, 4]
(4, 5]
(5, 6]
(6, 7]
(7, 8]
(8, 9]
(9, 10]
(10, 11]
(11, 12]
(12, 13]
(13, 14]
(14, 15]
(15, 16]
(16, 17]
(17, 18]
(18, 19]
(19, 21]
(21, 22]
(22, 23]
(23, 24]
(24, 25]
(25, 26]
(26, 27]
(27, 28]
(28, 29]
(29, 30]
(30, 31]
(31, 32]
(32, 33]
(33, 34]
(34, 35]
(35, 36]
(36, 37]
(37, 38]
(38, 39]
(39, 40]
> 40

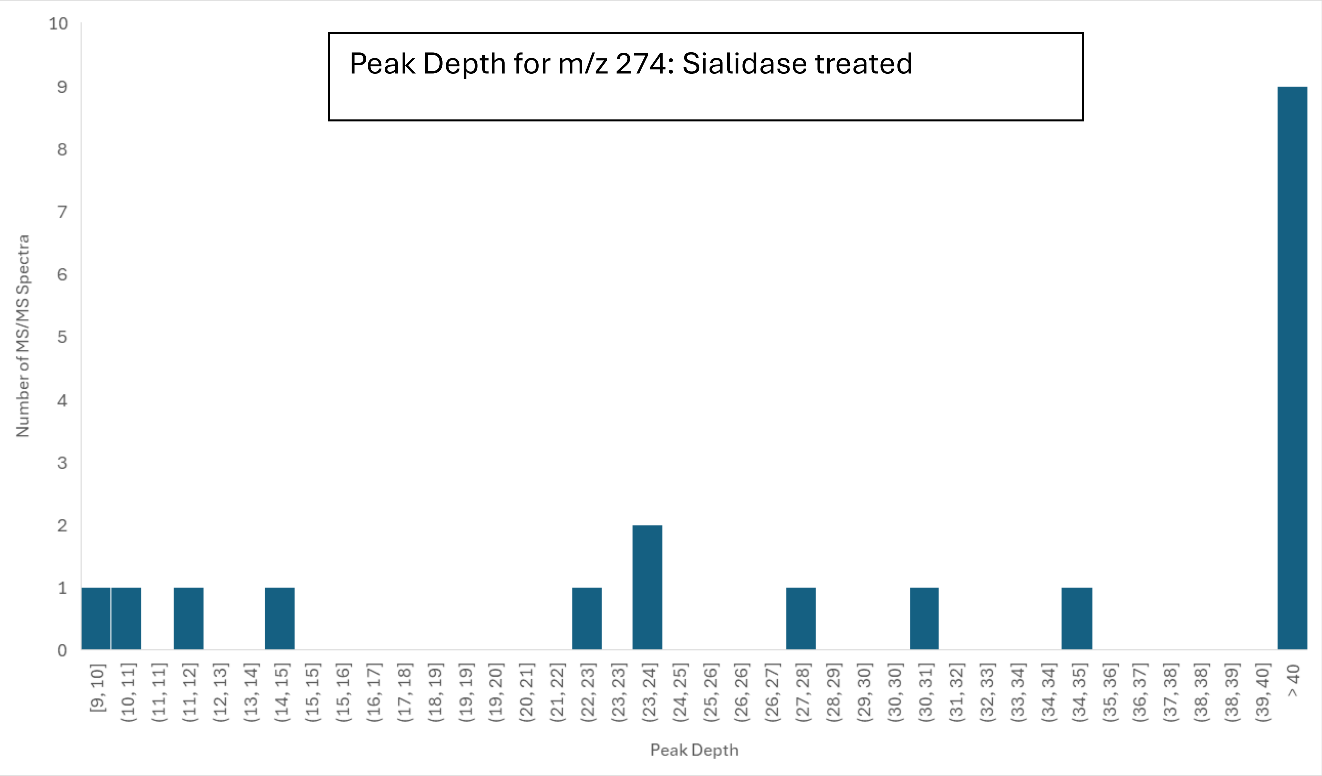
Peak Depth for m/z 274: Sialidase treated
Number of MS/MS Spectra
Peak Depth
0
1
2
3
4
5
6
7
8
9
10
[9, 10]
(10, 11]
(11, 11]
(11, 12]
(12, 13]
(13, 14]
(14, 15]
(15, 15]
(15, 16]
(16, 17]
(17, 18]
(18, 19]
(19, 19]
(19, 20]
(20, 21]
(21, 22]
(22, 23]
(23, 23]
(23, 24]
(24, 25]
(25, 26]
(26, 26]
(26, 27]
(27, 28]
(28, 29]
(29, 30]
(30, 30]
(30, 31]
(31, 32]
(32, 33]
(33, 34]
(34, 34]
(34, 35]
(35, 36]
(36, 37]
(37, 38]
(38, 38]
(38, 39]
(39, 40]
> 40

**OxoSignal:** The oxonium ion signal file displays the intensity values of each ion. Like the peak depth file, there is one column per ion, so we can compare the difference in intensities of certain ions either within a file or across different files. Here I will examine the difference in intensity of one of the sialic acid ions before sialidase and after sialidase (box plots below). Using the TotalOxoSignal column, you could also determine what fraction of the total signal each oxonium ion accounts for.

There is a summary column towards the end of this file that shows the fraction of the total ion current that was provided by oxonium ions (OxoTICfraction). This information is also used in the likely glycopeptide spectrum determination based on the TIC fraction setting from Step 5 of the GlyCounter User Interface section.  Both the peak depth and oxonium signal files show all the information that goes into the LikelyGlycoSpectrum determination and will also show true or false values in the final column.

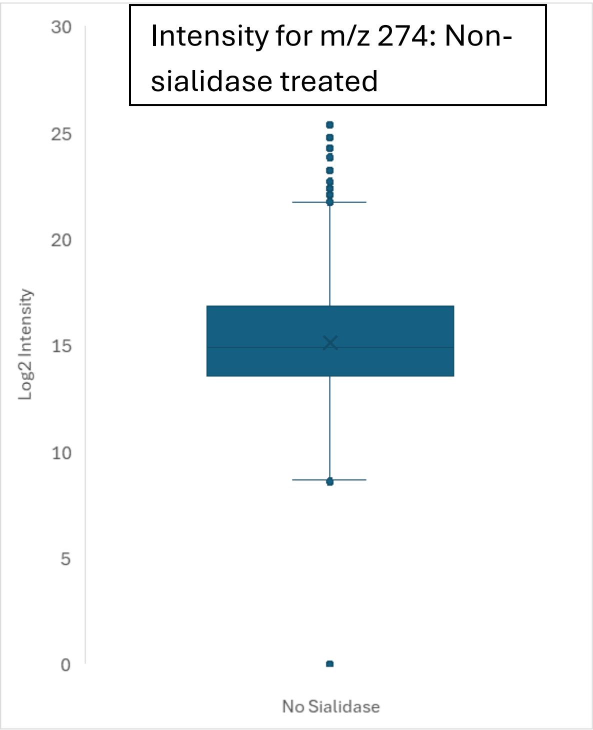


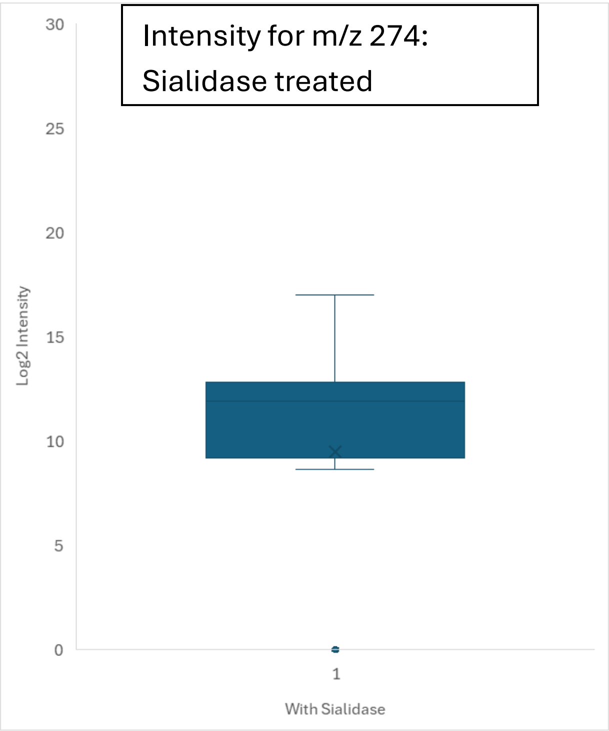


**Summary:** The GlyCounter summary file contains overarching information about the oxonium content. This file is great for a quick verification that a procedure worked correctly (for example, a high percentage of Likely Glyco spectra in a glyco-enriched sample). It also works well for comparing multiple runs. For this example, I looked at the percentage of spectra containing each of the sialic acid ions before and after sialidase treatment (bar charts below).

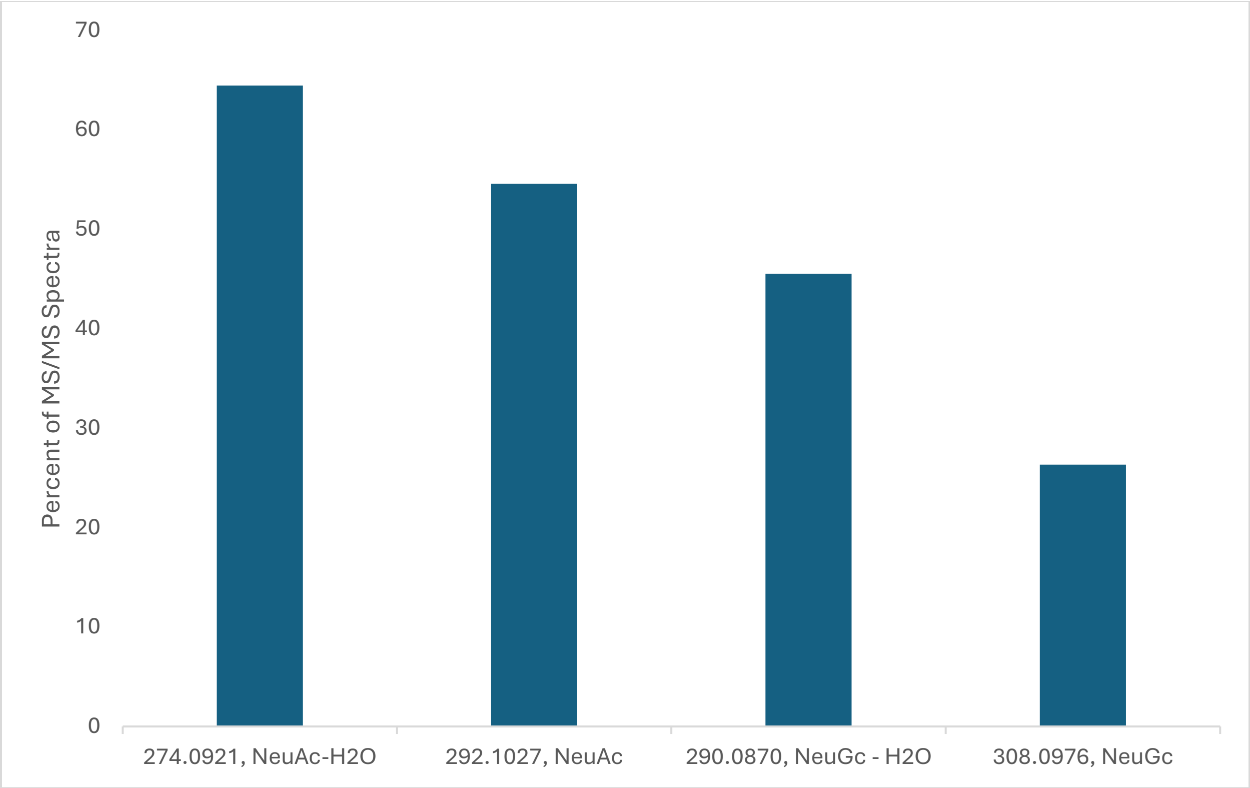

70
60
50
40
30
20
10
0
Percent of MS/MS Spectra
274.0921, NeuAc-H2O
292.1027, NeuAc
290.0870, NeuGc - H2O
308.0976, NeuGc


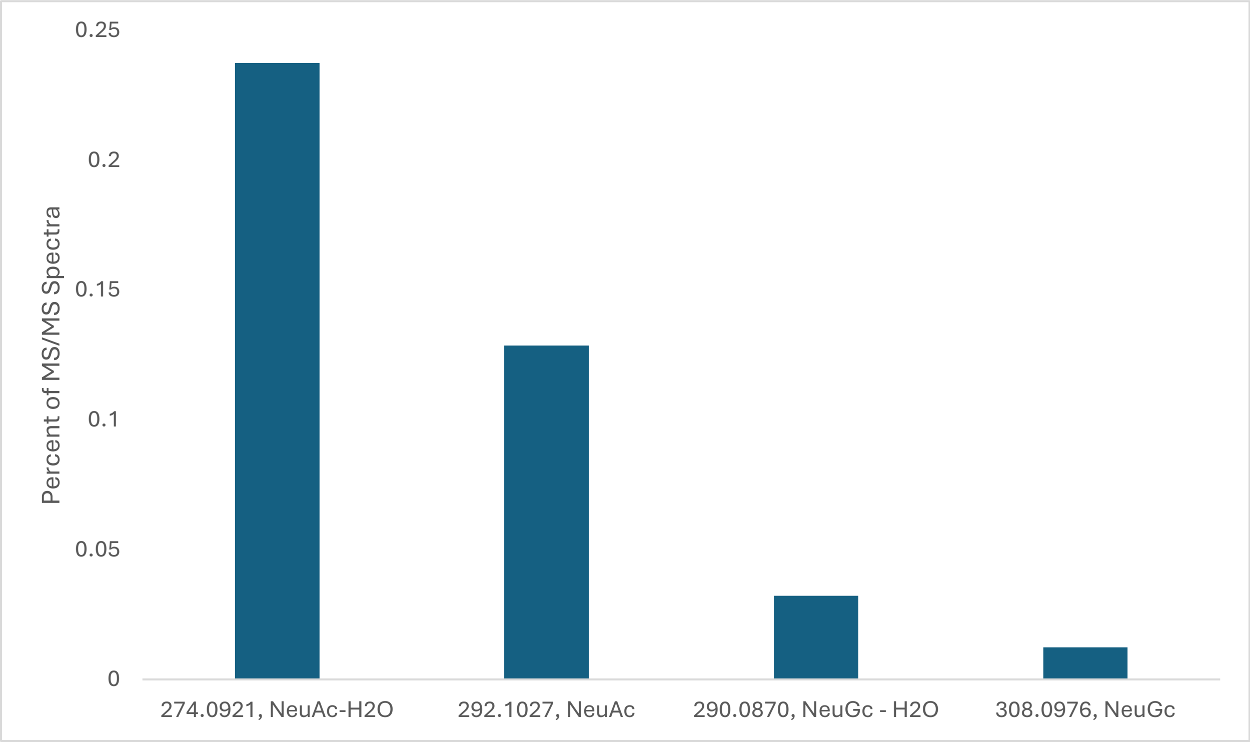

0.25
0.2
0.15
0.1
0.05
0
Percent of MS/MS Spectra
274.0921, NeuAc-H2O
292.1027, NeuAc
290.0870, NeuGc - H2O
308.0976, NeuGc

#### Ynaught

Compatibility for MS data files is the same for YNaught as GlyCounter. See the MSConvert section of this tutorial.

Ynaught is used to find Y-ions that appear in spectra. To do this, an identification file is required. MSFraggerGlyco search files from FragPipe are automatically compatible (see FragPipe documentation for how to run). Files searched with other algorithms and programs need to be formatted to match the sample Ynaught glycopeptide ID file found in the GitHub repo.

Step 1: Use the browse button next to the identification file upload (1) to select an identification file. See above for requirements.

Step 2: Upload a raw file using the browse button next to the raw file upload box (2).

Step 3: Select output directory by pressing the browse button next to the output directory field (3). By default, output files will be placed in the same location as the MS data file that was uploaded.

Step 4: Choose Y-ions. Y-ions labeled “Pep + monosaccharide(s)” build up from the peptide backbone. These ions are divided into categories corresponding to the type of glycan they most likely build, including N-glycans, core fucosylated glycans, and O-glycans. The glycan neutral losses category shows glycan losses from the complete glycopeptide. These are mostly ambiguous to glycan type. The Check All Ions or Check Common Ions buttons (4) can be used to select many ions at once. Custom additions or subtractions can be uploaded using the browse buttons at the bottom of the program (5).

Step 5: Fragment settings. Fragment ion tolerance can be adjusted in either ppm (default) or Dalton (by clicking the check box labeled Da) (6). The signal-to-noise (S/N) requirement for each peak can also be adjusted, as well as an intensity threshold (7). If noise data is available in the file (Thermo Orbitrap files) S/N will be used. Otherwise, the program will use the intensity threshold.

Step 6: Isotopes and charge states. Y-ions have isotope patterns that can help to verify that the peak being identified is not noise. Check the box next to the additional isotope(s) that Ynaught should try to find in the spectrum (8). Y-ions can have charge states of 1 up to the precursor charge state. All possible charge states can be searched for by leaving the default of 1 to P (where P is precursor charge) but specific charge states can be specified as well (9).  For more than one charge state, data will either be condensed into one column, or a column will be created for every possible charge state depending on the user selection

(10). Condensing data allows for smaller files and a cleaner view. Separate columns allow for the intensity of each charge state to be compared.

Step 7: Output IPSA annotations (11). Provides an additional output with ion, m/z, and mass difference columns. This is useful for spectral annotation.

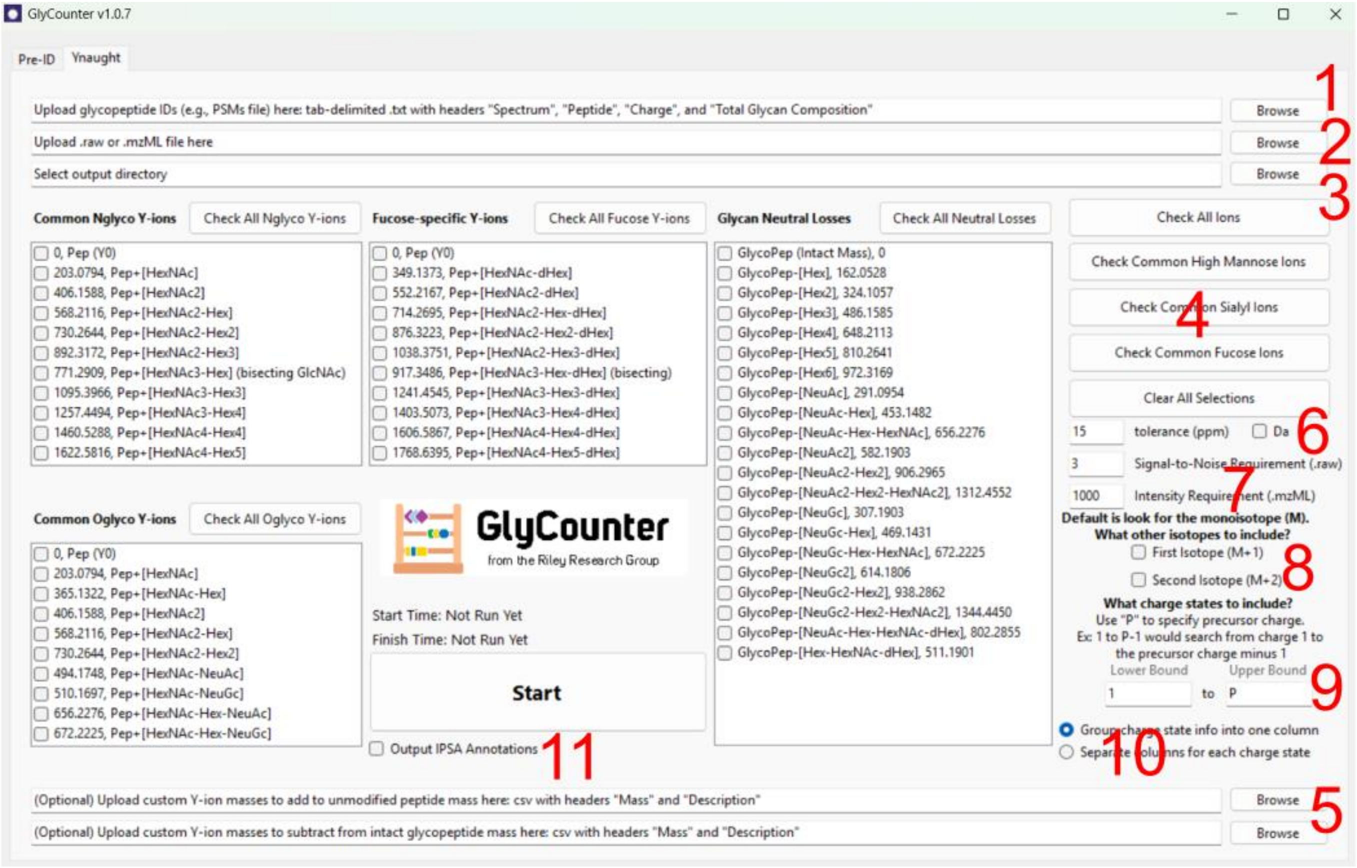


Ynaught outputs can be processed similarly to GlyCounter outputs. The Y-ion signal file contains intensities of the Y-ions (summed if "Group charge state info into one column" is checked). The Y-ion summary file has percentages of spectra that contain each Y-ion.
